# Probability Distribution for Rare Neutral Mutations in Cancers and Application to Dynamic Precision Medicine of Cancer

**DOI:** 10.1101/2025.10.10.681641

**Authors:** Wei He, Matthew D. McCoy, Chen-Hsiang Yeang, Rebecca B. Riggins, Robert A. Beckman

## Abstract

Cancers exhibit substantial genetic diversity between individual cells. Previous work shows greater overall diversity that also increases more quickly during a patient’s clinical course than heretofore expected. Rare subclones will harbor pre-existing resistance to any single agent and may cause medium to late term relapse, which may evolve further variants that have simultaneous resistance to non-cross resistant therapies. In this work, we present a probability distribution function (PDF) of the variant allele fraction (VAF), or prevalence, of a rare subclone, derived from previous evolutionary theory. We show that current clinical sequencing protocols fail to detect the vast majority of these rare subclones and that by the time of detection, simultaneous multiple resistance may already evolve. We apply the PDF to simulation of dynamic precision medicine (DPM), an evolutionary guided precision medicine paradigm that attempts to proactively eliminate singly-resistant subclones before they evolve multiple resistance, with significant potential to extend survival. We show the simulated benefit of DPM with perfect information is hampered by an inability to detect rare subclones if they are assumed to be absent when undetectable. However, the DPM benefit is restored if the PDF is used to calculate the likelihood of the subclone being present below the level of detection and incorporated into the DPM simulation and therapy recommendations in a probabilistic fashion. Two other common statistical distributions are less effective. This theoretical advance facilitates DPM and potentially other evolutionary guided approaches to precision cancer medicine in spite of the limitations of clinical sequencing.

**Significance Statement:** Cancers contain many cells, each genetically unique. These variations can include pre-existing resistance to therapy, enabling relapse. DNA sequencing cannot detect minority cellular populations (subclones) below a certain size, by which time they may have already evolved simultaneous resistance to multiple therapies. We present a mathematical approach that assigns a risk that a subclone is present, and at what prevalence in the cellular population, even when undetected. We simulate applying this approach to dynamic precision medicine (DPM), which attempts to proactively eliminate singly resistant cells before they become resistant to multiple therapies. Using this probability distribution, we can retain the benefit of DPM even when most rare subclones are undetectable, in contrast to just assuming undetected subclones are absent.

## Introduction

Cancers exhibit significant genetic diversity among individual cells. Previous work using colorectal cancer diagnostic biopsy specimens (1, 2) has shown a mutation rate of 7.1 × 10^−7^ per base per new cell added to a growing cancer, several orders of magnitude higher than previously believed (3). This work employed duplex sequencing, a technique that provides exceptional sensitivity and specificity for mutation detection (4–6), with sufficient accuracy to sequence at a depth of 1 million in principle Moreover, the theoretical approach demonstrated that the infinite sites assumption (7), equivalent to the assumption that a given mutation is newly formed only in one cell at a time in a single doubling of a cell population, did not apply when the number of cells in the total cancer exceeds the reciprocal of the mutation rate, in this case several orders of magnitude below the number of cells in a single radiologically detectable lesion (1, 2). Previous models (8–10) predicted that the variant allele frequency (VAF) or prevalence of a mutation in the cancer cell population would be inversely proportional to the number of cells, whereas the new model (1,2) predicts that the VAF will level off at the mutation rate, consistent with the fact that physico-chemical properties of the enzymatic systems within living cells have accuracy limits (11). As we have shown, the predictions of our model begin to diverge from other models when a systemic cancer has a total volume of 1 mm^3^ throughout the body. Yet, this divergence is invisible unless sequencing depths reach the order of 1 million (1, 2). Based on this model, a far greater increase in genetic diversity is predicted as the number of total cells in the cancer increases, compared to earlier models. Moreover, any single resistance mechanism would be present in one or more cells in any lesion large enough to be diagnosable, unless that resistance mutation is highly selected against. Indeed, in a one cm^3^ lesion containing 10^9^ cells, we expect that 700 cells with any single mutation would be added to the lesion with every population doubling by mutation from sensitive cells, making stochastic elimination of this heterogeneity extremely unlikely (1, 2).

Based on these experimental and theoretical results, we extend the theory further to examine predicted evolution of rare subclones as a potential source of therapy resistance. We derive a probability distribution function (VAF-PDF) expressing the probability that a neutral mutation will be present at a given variant allele fraction (VAF) at diagnosis. For diploid cells, we assume that the mutation is dominant and on a single chromosome, so that the subclonal PDF is twice the VAF-PDF. Clinical sequencing protocols rarely detect subclones below 1% VAF (12–14). We find that the vast majority of neutral genetic diversity in a cancer at diagnosis is well below the level of detection afforded by clinical sequencing protocols. Rare subclones may take longer to grow out and thus be the cause of medium to late term relapses. Precision medicine has led to considerable patient benefit by targeting subclones with specific drugs (15–18). However, due to the development of therapy resistance, relapse is a frequent problem, meaning cures are elusive. Prevention of medium to late term relapses occurring in months to years will be an increasing focus of oncology as short-term results reflecting initial responses continue to improve. The time scales of and definitions of medium and late relapses will vary depending on the cancer type and individual patient evolutionary dynamics. Genetic diversity is one of several causes of therapy resistance, and perhaps the major cause of irreversible resistance, as compared to reversible resistance due to gene expression changes in the absence of genetic change (19). Genetic subclonal diversity is divided into mutations that provide a selective fitness advantage to cancer cells, commonly referred to as *drivers*, and mutations that are neutral with respect to their effect on fitness, or *passengers*. Driver mutations are expected to have a high variant allele frequency (VAF) due to evolutionary selection. Our previous work focused on the evolution of passengers, and considered only mutations with VAF of 0.1 (10%) or below (1, 2). In many cases, the effect of a single mutation on fitness may be dependent on the context provided by the microenvironment near the cell, as well as other mutations in that cell’s genome.

Importantly, once therapy begins, the drivers targeted by therapy may have a negative effect on fitness, due to therapy-induced cytostasis or cytotoxicity in cells that harbor them. Alternate drivers or further modified forms of the original drivers may also appear. We term these *driver resistance mutations*. In contrast, some former passenger mutations that did not provide a fitness advantage during initial carcinogenesis may provide a source of therapy resistance, and therefore provide a selective fitness advantage for a subclone only in the presence of therapy. We term these *neutral resistance mutations*. Rare subclones harboring these neutral resistance mutations (former passenger mutations) may take longer to grow out and thus be the cause of medium to late term relapses Additionally, these *neutral resistance mutations* may in principle occur from unanticipated loci, including intronic segments affecting gene expression, and thus the percentage of the genome that may be susceptible to them may be large.

Current precision medicine (CPM) targets therapy to the molecular drivers of the largest subclones in a cancer, repeating the process at relapse, with progressively decreasing clinical benefits at each iteration. Evolutionary guided precision medicine (EGPM) paradigms (20–28) more explicitly embrace the subclonal genetic heterogeneity and evolutionary dynamics of cancer in an effort to delay or prevent resistance. The inability to detect rare subclones may affect the utility of these methods.

Dynamic precision medicine (DPM) (20) is one of several early EGPM approaches. It is currently in early stages of preclinical testing. Simulations suggest that it can provide significant improvements in survival and cure rates across a broad range of clinical presentations by providing a personalized sequence of therapies of interest (20, 29) based on measured parameters relevant to heterogeneity and dynamics, including subclonal growth rates, drug sensitivities, and mutation rates. DPM does not discover or suggest new therapies, but rather recommends the optimal therapy sequence for a given patient when a menu of non-cross resistant “drugs” is provided. Each ‘drug” may itself be a combination of therapies, and there may be synergy and/or cross resistance within each “drug”. In contrast to CPM, which treats with the same drug until relapse or worsening of the cancer, DPM suggests frequent proactive adaptations of therapy (“moves”), but the first two moves provide most of the benefit, enhancing the feasibility and cost-effectiveness (30). Simulations suggest that some virtual patients will benefit more than others, and a method for selecting patients who will benefit has been published (30). The model has been extended to include reversible non-genetic mechanisms (DPM-J) (19). Notably, the model does not assume that all subclones have the same mutation rate, but that some subclones will be “hypermutators” (31) based on random mutations in the cellular machinery that protects the integrity of the genome. These hypermutator subclones may be particularly dangerous due to their rapid evolution. Assumptions underlying DPM have been previously presented (20, 30).

Simulations of DPM show that very rare subclones, as low as VAF 1:100,000, may alter the optimal sequence of therapies (20, 32). If the vast majority of subclones are indeed exceedingly rare, below the level of clinical detection, the question arises as to whether the inability to detect the subclones will reduce or even eliminate the potential benefits of DPM in preclinical and/or clinical testing. To test the potential application of the VAF-PDF, we simulate the benefit of DPM compared to CPM with decreasing levels of sensitivity for subclone detection in two ways. In one approach, if no resistance is detected at diagnosis, we assume it is absent until the resistant subclone grows to a detectable size and is detected. In an alternate approach, if resistance is not detected, we assume that it is present, based on our prior work (1, 2), but at a subclonal prevalence and corresponding VAF below the limit of detection (LOD). Using the VAF-PDF, we calculate the probability that resistance is present at varying levels below the LOD at diagnosis, and then it evolves according to the DPM equations during therapy. We then weigh the DPM recommendations from different VAF levels by their probabilities, resulting in a personalized probability-weighted optimal therapy sequence. We compare the robustness of DPM to limitations in subclone detection sensitivity by these two approaches, as well as using other commonly used statistical distributions not based on evolutionary theory to approximate VAF if a subclone is undetected.

We find that the benefit of DPM is reduced (but not eliminated) when the sensitivity of subclone detection is decreased and resistant subclones are not considered until they become detectable. However, using the VAF-PDF as described above restores the benefit of DPM to close to its original benefit level. Moreover, the VAF-PDF is more effective for this purpose than other common statistical distributions. Additionally, we explore potential methods and approaches to address inconsistences between simulated results and the actual cancer cell population observed by clinicians. These insights could be valuable when applying model suggested treatments in clinical practice.

## Results

### Defining the Likelihood of VAFs at Neutral Sites in Cancers at Diagnosis

Current sequencing depths reveal only a fraction of the underlying genetic diversity. Our previous work demonstrated that genetic diversity among individual cancer cells is far greater than previously appreciated and increases more rapidly over the course of a patient’s clinical progression than earlier anticipated (1, 2). Rare subclones carrying pre-existing resistance to a single therapeutic agent, often responsible for relapse in the medium to late stages, typically remain undetectable by standard clinical sequencing until they expand to a detectable size in a frank relapse, by which point they are large enough to contain significant sub-subclonal diversity. Therefore, it is essential to first derive or define a PDF for the VAF (VAF-PDF), or subclone prevalence, based on established evolutionary theory. As derived in the *Materials and Methods* section, equation 9 defines the PDF for the VAF and equation 11 computes the fraction of neutral genome sites across different ranges of VAF.

Table 1 is produced by using the cumulative probability for the VAF *ρ* to be within a range between *ρ*_*min*_ and *ρ*_*max*_. In Table 1, the probabilities assume the VAF *ρ* can range from *ρ*_*min*_ or, the intrinsic effective mutation rate per new cell added, to *ρ*_*m*_ or 0.5 (heterozygous mutations in all cells). Fig. 1 further illustrates how the proportion of neutral genomic sites varies across different VAF ranges, with each line color representing a different starting *ρ*_*min*_ and the x-axis indicating the corresponding *ρ*_*max*_. Our results confirm that the vast majority of neutral mutations are predicted to reside in rare subclones that are undetectable by current clinical sequencing methods. Table 1 shows that only a very small fraction of mutations occurs at allele frequencies of 10^−2^ or higher, a common detection threshold for conventional DNA sequencing (33–35). Specifically, in the second to last row of Table 1, the probability of a mutation falling within this range is approximately 0.000065, also indicated by the cyan and yellow color lines in Fig. 1. In contrast, 93% of neutral variants have a VAF of 10^−5^ or lower, as shown by summing the second column of the first two rows in Table 1 or by the black curve in Fig. 1 at *ρ*_*m*_ = 10^−5^. Moreover, over 99% of variants have a VAF of 10^−4^ or lower, based on the sum of the second column in the first four rows of Table 1 or the black curve in Fig. 1 at *ρ*_*m*_ = 10^−4^. When = 7.1 × 10^−7^, the average VAF, as calculated using equation 12 (*Materials and Methods*), is 9.6 × 10^−6^. This corresponds to an average subclone size of approximately 9400 cells, assuming a total cancer burden of 10^9^ cells (*Materials and Methods*).

**Table 1.** Fraction of neutral genome sites at different ranges of VAF in an untreated cancer.

| $\rho_{min}$ | $\rho_{max}$ | Fraction of mutated genome sites | Total mutated genome sites |
| --- | --- | --- | --- |
| $7.1 \times 10^{-7}$ | $1.12 \times 10^{-6}$ | 0.366 | $1.1 \times 10^9$ |
| $1.12 \times 10^{-6}$ | $1 \times 10^{-5}$ | 0.563 | $1.69 \times 10^9$ |
| $1 \times 10^{-5}$ | $1 \times 10^{-4}$ | 0.064 | $1.92 \times 10^8$ |
| $1 \times 10^{-4}$ | $1 \times 10^{-3}$ | 0.0064 | $1.92 \times 10^7$ |
| $1 \times 10^{-3}$ | $1 \times 10^{-2}$ | 0.00064 | $1.92 \times 10^6$ |
| $1 \times 10^{-2}$ | $1 \times 10^{-1}$ | 0.000065 | $1.95 \times 10^5$ |
| $1 \times 10^{-1}$ | 1 | 0.0000071 | $2.13 \times 10^4$ |

Each range is from $\rho_{min}$ to $\rho_{max}$ . The top rows represent the rarest mutations. The number three billion is used as the total nucleotide base pairs in the human genome.

**Figure 1.**
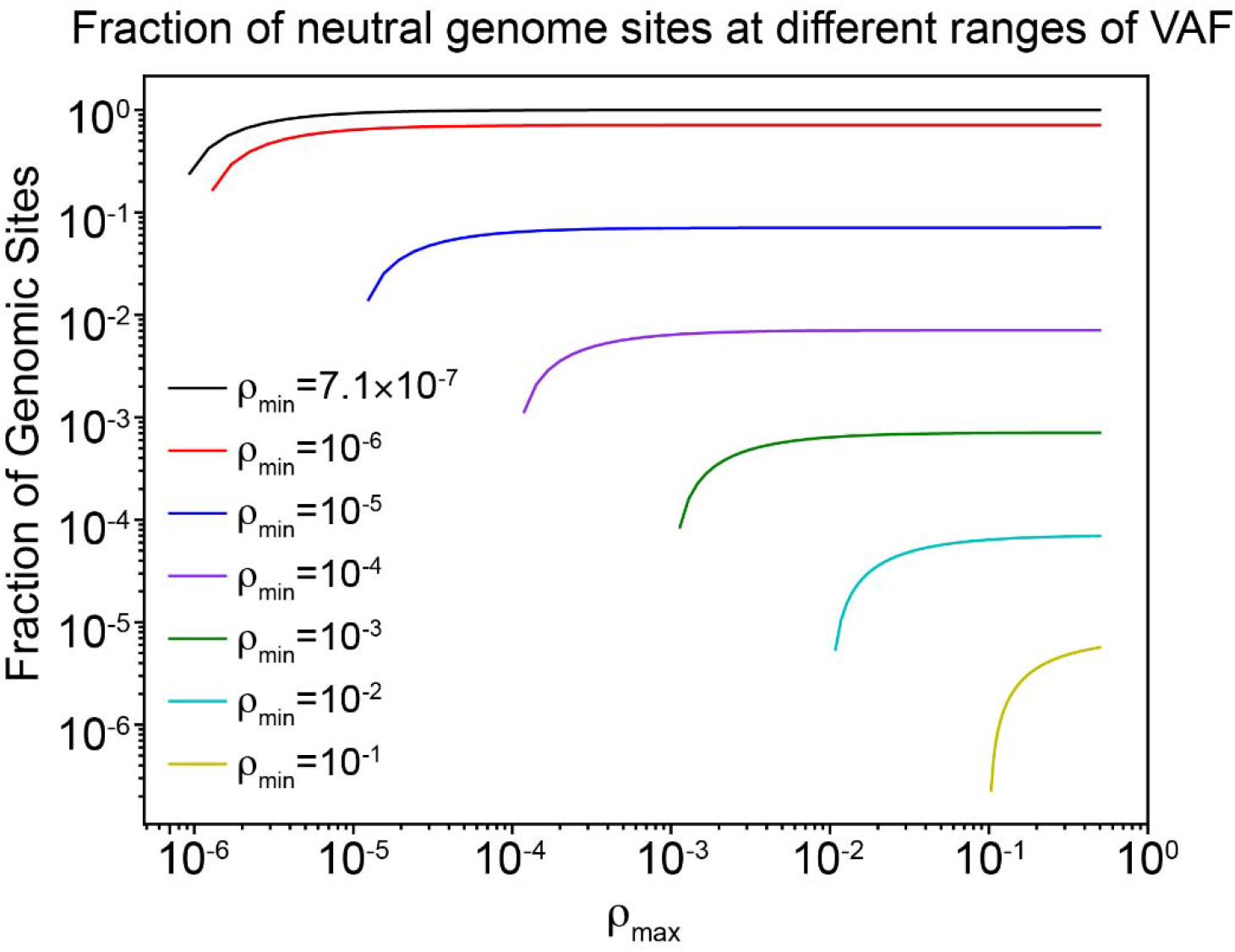
Proportion of neutral genomic sites across varying VAF ranges. Values are calculated using equation 11, with *ρ*_*min*_greater than or equal to the effective mutation rate, determined to be 7.1×10−□in colorectal cancer art diagnosis (1) and *ρ*_*max*_ less than or equal to 0.5, corresponding to a heterozygous mutation in every cell, assuming a diploid population, using different combinations of *ρ*_*min*_ and *ρ*_*max*_. In a practical application, *ρ*_*min*_will be equal to the limit of detection (LOD), typically 10^−1^ or 10^−2^ with today’s clinical sequencing protocols.

After deriving this key probability density function for the VAF (VAF-PDF), we incorporate it into the CPM and DPM simulations to address model input parameter mis-specification due to imperfect detection of rare subclones. We account for the possibility that subclones may exist below the detection threshold. By using the VAF-PDF to estimate the likelihood of these undetectable subclones and integrating this uncertainty probabilistically into DPM simulations and treatment recommendations, we are able to recover much of the lost benefit due to undetected subclones.

### Mis-specification adversely affects survival curves of DPM treatment strategy

Fig. 2*A* and *C* show the Kaplan-Meier curves of CPM and DPM treatment strategies without mis-specification of subclonal prevalence input parameters for virtual patient sets one and two, respectively. Each virtual patient corresponds to a different set of input parameters at diagnosis: the prevalence of drug sensitive subclones and subclones singly resistant to drug 1 and drug 2, respectively, and the net growth rates, mutation rates, and drugs sensitivities of all subclones. Virtual patient set one is the patient set used in the original DPM simulation, in which singly resistant subclone prevalence is varied across a broad range evenly spaced on a logarithmic scale (20). For virtual patient set two, we sample the VAF-PDF (multiplied by 2) to determine the prevalence of the singly resistant subclones. We can see that for both virtual patient sets, DPM (blue) curves are significantly superior to CPM (red). DPM demonstrates a significantly higher median survival time compared to CPM, with 386 days versus 220 days for virtual patient set one and 816 days versus 273 days for virtual patient set two, respectively. Additionally, the hazard ratios of DPM relative to CPM are 0.518 and 0.429 for virtual patient set one and two, respectively. The new findings from virtual patient set two are consistent with our previous results which used virtual patient set one, demonstrating that the DPM strategy outperforms the current precision medicine strategy (20). These results emphasize the importance of preventing the emergence of doubly resistant subclones to both drugs.

**Figure 2.**
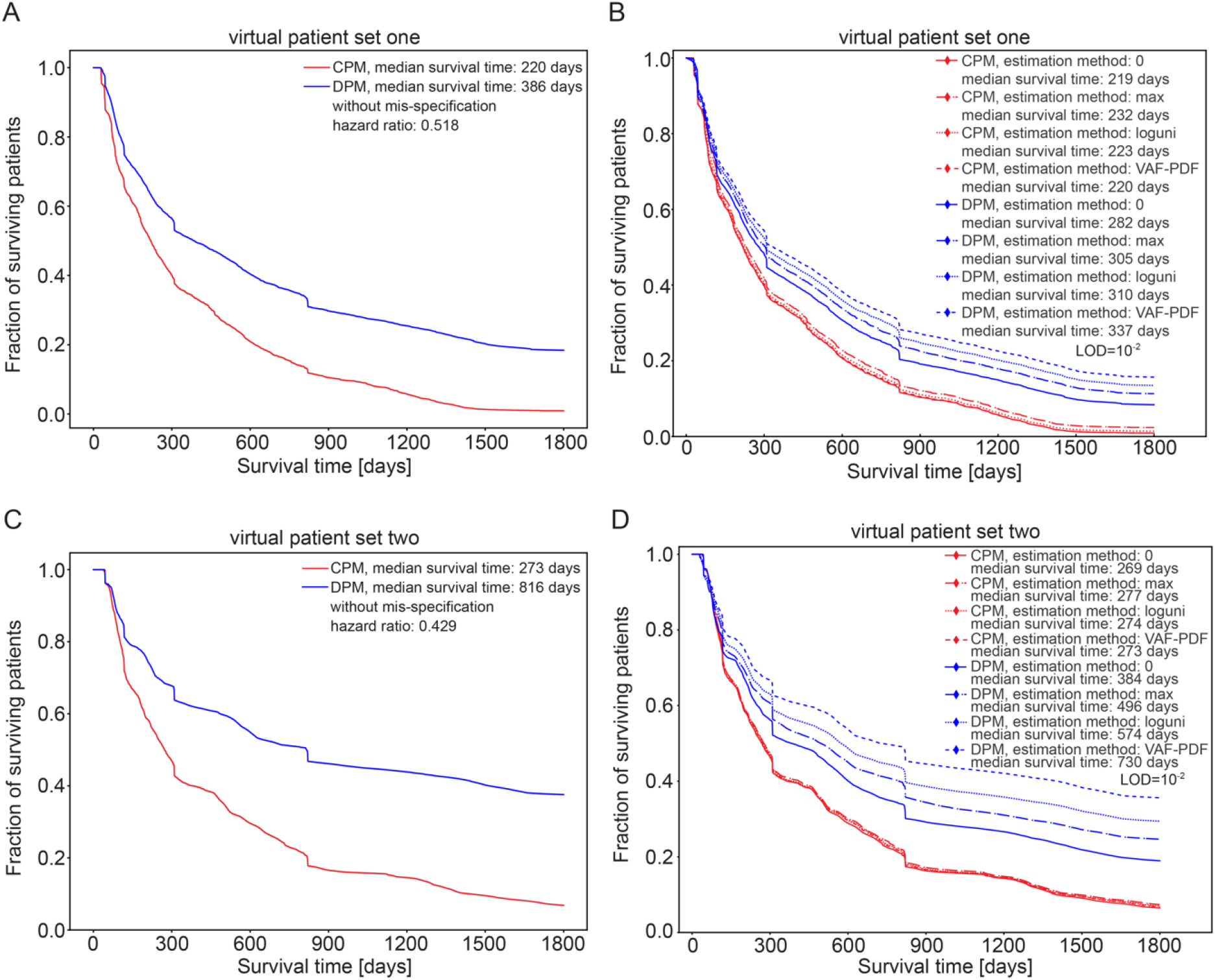
Kaplan-Meier curves comparing CPM and DPM treatment strategies under mis-specification with an LOD of 10^−2^ and without mis-specification for virtual patient set one and set two. (*A*) Kaplan-Meier curves illustrating CPM and DPM outcomes for virtual patient set one without mis-specification. (*B*) Kaplan-Meier curves illustrating CPM and DPM outcomes for virtual patient set one with mis-specification under different estimation methods, with LOD equal to 10^−2^. (*C*) Same as (*A*) but for virtual patient set two. (*D*) Same as (*B*) but for virtual patient set two. Virtual patient set one is the patient set used in the original DPM simulation (20). For virtual patient set two, we sample the VAF-PDF (multiplied by 2) to determine the prevalence of the singly resistant subclones.

Fig. 2*B* and *D* present the Kaplan-Meier curves of CPM and DPM treatment strategies under mis-specification, using different estimation methods to estimate subclonal prevalence below the limit of detection (LOD) with an LOD of 10^−2^ for virtual patient sets one and two, respectively. The LOD of 10^−2^ is used for illustration because it reflects current clinical sequencing practices. The Kaplan-Meier curves for all combinations of estimation methods and LODs for virtual patient set one and set two are shown in *SI Appendix* Fig. S1 and S2. The estimation methods when no subclones are detected are method 0 (assume 0 when undetected), method max (assume subclonal prevalence just below the limit of detection), method loguni (a log-uniform distribution, assuming a logarithmic probability distribution) and method VAF-PDF (assume the VAF-PDF multiplied by 2 for the subclonal prevalence). In Fig. 2*B*, we can see that mis-specification of subclonal prevalence has little impact of the CPM strategy across all methods, with median survival times nearly identical to the scenario without mis-specification at 220 days for virtual patient set one. However, for the DPM strategy which aims to prevent the emergence of subclonal resistance, all methods of estimation regarding the subclones lead to a reduction in the median survival time. The median survival times for estimation method 0, max, loguni and VAF-PDF are 282 days, 305 days, 337 days and 336 days respectively, all lower than 381 days in the scenario without mis-specification. The two extreme methods dealing with the limit of detection, either setting the subclone percentage to 0 or to the maximum, result in relatively lower survival times compared to the loguni and VAF-PDF methods. As the actual subclone percentages lie between 0 and the maximum, assigning their values to these extremes is simplistic and diminishes the effectiveness of DPM. Moreover, neglecting the existence of rare subclones by setting their values to 0 leads to the lowest median survival time. This approach directly conflicts with DPM’s goal of preventing resistant subclones, significantly undermining its benefits. The other two approaches are sampling the subclone prevalence probabilistically between their extreme values. Using a log-uniform distribution, we assume that the logarithm of the subclone percentages has equally likely outcomes, without favoring the lower or higher percentages, while the VAF-PDF method relies on the VAF-PDF derived herein using evolutionary theory (see *Materials and Methods*). The two sampling approaches achieve better results than the extreme methods, with the evolutionary derived VAF-PDF showing a longer median survival time. Given that in virtual patient set one, the VAFs were initially set to a log-uniform distribution, the superior performance of the theoretically derived PDF is a striking result. This suggests that even if the true VAF-PDF diverges from theory at diagnosis, it will approach the theoretical VAF-PDF with further evolution, subject only to the elimination of other subclones by therapy.

Fig. 2*D* demonstrates the Kaplan-Meier curves for the virtual patient set two. Mis-specification has minimal impact on the CPM strategy across all methods, with median survival times nearly identical to the perfect scenario without mis-specification. Similarly, all estimation methods on subclone percentages result in reduced median survival times for the DPM strategy. But unlike virtual patient set one, the distribution of the subclones percentage in virtual patient set two follows the probability density function described in equation 9. In this case, sampling the subclone percentage distribution using the same distribution as in the virtual patient set significantly mitigates the impact of mis-specification. This is reflected in the longer median survival time of 726 days with the VAF-PDF method, compared to 384 days, 496 days, and 574 days with the 0, max, and loguni methods, respectively.

### Examine the impact of mis-specification on clinical metrics under different LODs

Fig. 3 shows the hazard ratio of DPM:CPM as a function of the LOD for virtual patient sets 1 and 2. Results are consistent with the Kaplan-Meier curves in Fig. 2. Assuming 0 for the subclonal prevalence when undetected gives the worst results, and using the VAF-PDF derived from evolutionary theory gives the best results. In the case of virtual patient set 2, almost all of the benefit is preserved when using the VAF-PDF method. Median survival time, absolute hazards, and changes in case numbers showing that DPM performs significantly better than CPM are shown in *SI Appendix* Fig. S3. In addition, *SI Appendix* Fig. S4 presents cases from individual virtual patients under the LOD 10^−2^ scenario, which reflects current clinical sequencing practices. These examples demonstrate how mis-specification of subclonal prevalence can disrupt optimal sequencing recommendations and reduce survival.

**Figure 3.**
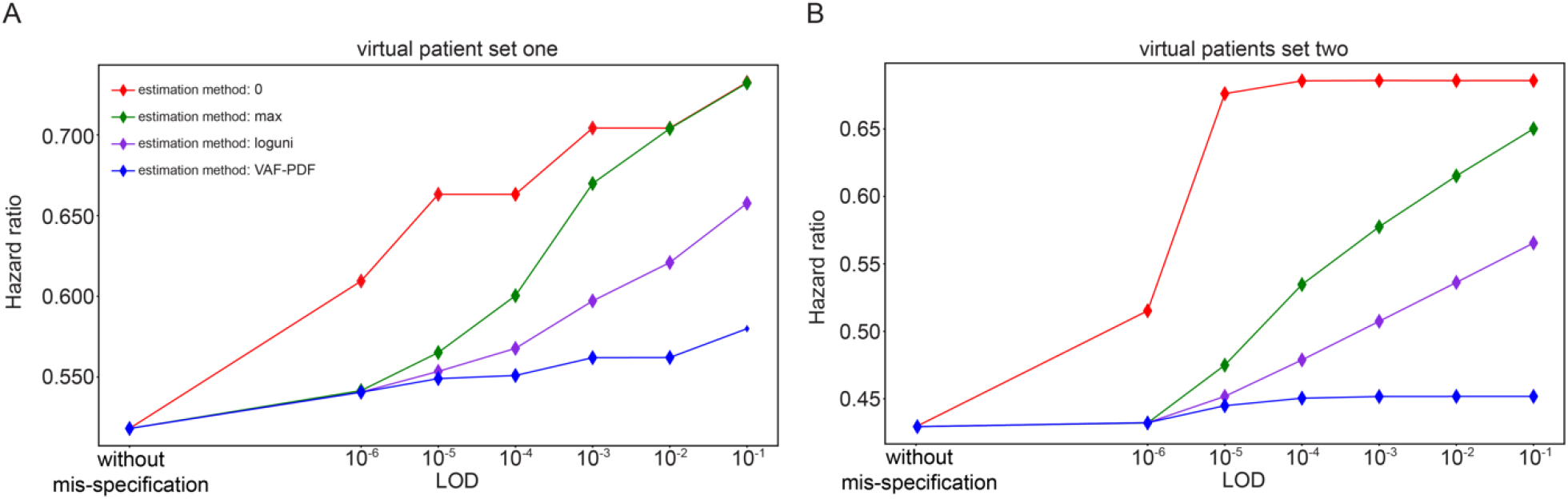
Comparison of hazard ratios under scenarios with and without mis-specifications and varying LODs for virtual patient set one and two. (*A*) Changes in hazard ratios of DPM relative to CPM across various estimation methods, with increasing LODs, starting from the scenario without mis-specification for virtual patient set one. (*B*) Same as (*A*) but for virtual patient set two. Virtual patient set one is the patient set used in the original DPM simulation (20). For virtual patient set two, we sample the VAF-PDF (multiplied by 2) to determine the prevalence of the singly resistant subclones.

## Discussion

We present an analysis of the evolution of cancer subclonal genetic heterogeneity for neutral passenger mutations. Our analysis suggests that current DNA sequencing technologies used to support precision oncology fail to detect most of this heterogeneity, which we hypothesize to be a contributor to therapeutic resistance and medium to late term relapses. We have derived a PDF for the VAF of rare subclones evolving neutrally before diagnosis. Specifically, while clinical sequencing protocols generally can detect subclones at VAF 10^−1^ higher, or on rare occasions 10^−2^ or higher, our analysis indicates that >99% of subclonal diversity is at VAF 10^−4^ or lower, and >93% is at VAF 10^−5^ or less. The mean VAF is on the order of 10^−5^, or a mean subclonal size of about 10^4^ cells per 1 cm^3^ lesion, a number that is likely too large to be consistent with elimination by stochastic population fluctuations.

We show by simulation that failure to detect rare subclones harboring drug resistance can hamper the benefit from the use of DPM, an evolutionary guided precision medicine paradigm. This effect is ameliorated if, instead of assuming a subclone is absent if undetected, the PDF is used to create a probability weighted prediction of the VAF of the subclone below the detection limit, and DPM then proceeds in a probability weighted fashion. Survival benefit in the simulations is specific to this PDF and not to other common statistical distributions, suggesting the potential for significant clinical utility specific to this PDF. Laboratory and clinical validation of DPM is in early stages.

These findings build on prior experimental and theoretical work (1, 2), which suggests that cells that are singly resistant to any therapeutic agent are already present in any cancer with more than 7 cubic millimeters of total volume across all observed and microscopic lesions (ca 2.37 mm diameter for a spherical lesion), unless there is no locus, mutation of which can lead to resistance without severe impact on cellular fitness. The experimental work involved ultradeep sequencing of 13 kb of DNA from patients with colorectal cancer at diagnosis, at various depth up to 20,000 (1), using duplex sequencing (4–6) a technique with very high accuracy and sensitivity. The mathematical model predicted the increasing detection of rarer SNVs with increasing depth, with correlation coefficients of 0.95 or higher. An analysis of individual bases showed that the base-to-base variance in SNV rates was consistent with a Poisson process rather than a Luria Delbruck process (1). These findings then formed the basis for the development of the PDF described herein.

Other key assumptions then fully determined the model. We modeled evolution as a branching process such that earlier mutations would have higher VAFs, and that these sites would be unavailable for later mutations, since their VAF could be well approximated by the earliest mutation at the given site. Reversions were assumed to occur only in a minority of cells, and the cancer as a whole was assumed to experience exponential growth during carcinogenesis from formation of the founder cell to diagnosis.

Bulk sequencing at a depth of 1 million or more would be required to comprehensively capture this heterogeneity. Duplex sequencing (4–6) and nanoseq (36) have sufficient accuracy to enable sequencing at this depth without confounding by noise, but are prohibitively expensive at that depth. Single cell sequencing does not circumvent this problem, as the sequencing of 1 million cells will result in multiple false reads in a subset of cells, and therefore detection is still limited by the noise level of the sequencing technology. Bioinformatic approaches involving the requirement for multiple confirmatory reads of a variant will reduce sensitivity. A full analysis of the consequences for sensitivity and specificity of subclonal detection is in preparation and will be published elsewhere. However, it seems clear that detection of the majority of mutant subclones will require lower cost technologies.

Our analysis (1, 2) differs from prior analyses in that it does not rely on the infinite sites assumption, which we showed to be inapplicable once the number of cells in the cancer begins to approach the reciprocal of the effective mutation rate per new cell added to the population, or about 1.5 million cells (1.5 mm^3^). As shown in Fig. 4, a much greater fraction of the heterogeneity is expected to be at low subclonal prevalence than in other models, increasing the importance of rare mutations compared to previous estimates. However, the figure also shows that our model, in contrast to other models, does not predict extremely rare subclones, and in fact states that the subclonal prevalence cannot be lower than the error rate of the DNA polymerase complexes for neutral mutations. Thus, a sequencing depth approximately equal to the reciprocal of the error rate of the polymerase complexes should be adequate, if it could be economically feasible in the future. Further, as described in *Materials and Methods*, the branching nature of evolution means variants that occur earlier in evolution have a higher subclonal prevalence. This means that by the time the lower limit of subclonal prevalence is reached, many of the loci have mutated earlier. Thus the mean subclonal prevalence is somewhat higher than the lower limit defined by polymerase accuracy.

**Figure 4.**
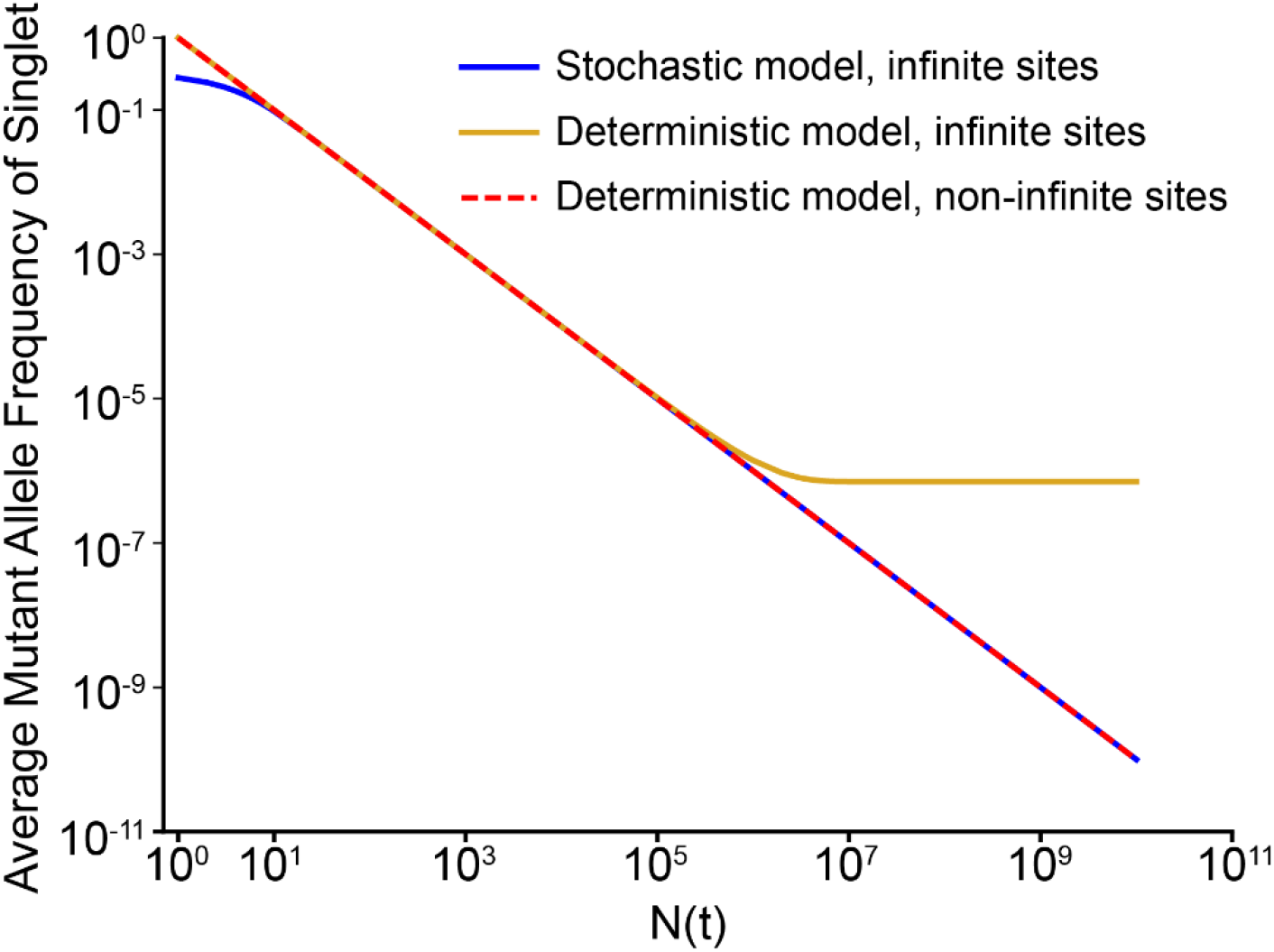
Comparison of model predictions. Average mutant allele frequency (MAF, mean VAF), is plotted against N(t), the number of cells at the time of mutation, for the stochastic model with the infinite sites assumption (9, 10) (blue), the deterministic model with the infinite sites assumption (8) (red), and without the infinite sites assumption (1, 2) (gold, equation 5 with *k* = 7.1 × 10^−7^) Sequencing to a depth D detects (on average) events formed when N(t) ≈ D. Duplex sequencing as done in (1) probes up to N(t) ≈ 20,000. The gold curve beyond this range is inferred from the equation 5 with *k* = 7.1 × 10^−7^. Within the directly observed range, stochastic models capture random fluctuations and are more accurate early in tumor growth, whereas deterministic models represent the average trajectory. Parameters are cell birth rate = 0.25/day, cell death rate = 0.18/day, effective mutation rate per base per new cell added to the tumor = 7.1 × 10^−7^. Adapted from references (1, 2).

The current analysis assumes that the mutation rate is constant in all cells, but in fact our earlier work (1, 2) implies at diagnosis there will be “hypermutator subclones” with random mutations in any of the large number of proteins responsible for maintaining genomic integrity, since all mutations will be present. These hypermutator subclones may then be further selected by therapy because they are more likely to harbor multiple resistance mutations simultaneously (31). This phenomenon may lead to higher effective mutations rates and more diversity after diagnosis, but would also raise the lower limit of subclonal prevalence, requiring somewhat less sequencing depth for characterization.

Insufficient sequencing depth is not the only limitation of the current exploration of cellular genetic heterogeneity within a single patient’s cancer. Current precision medicine practice often involves sequencing only the exome (1% of the genome, omitting promoters, enhancers, microRNA binding sites and other intronic loci that have important functions), or worse, a subset of exons already known to be important in cancer. This is again driven by economic considerations, but it tends to focus us on what we already know and narrows opportunities for further discovery. Enhanced sequencing of cancers will lead to the observation of many mutations of unknown significance. CRISPR resistance screens (37) as well as theoretical methods involving molecular dynamics simulations and artificial intelligence (38, 39) can begin to clarify some of these issues. Deeper sequencing may reveal new and important drug targets related to resistance to the current agents.

While our work implies the need for deeper exploration of genetic heterogeneity between individual cancer cells in a single patient, that is not the only source of heterogeneity and therapy resistance in cancer. Reversible resistance mechanisms at the transcriptomic, proteomic, and post-translational levels are known. We have begun to address some of these resistance mechanisms in our joint model of dynamic precision medicine, DPM-J, that considers both non-genetic and genetic resistance mechanisms (19). Interactions between cancer cells themselves as well as between cancer cells and the host microenvironment are also known to influence resistance. Of note, greater variability between lesions in response to therapy is seen in the context of immunotherapy compared to other therapies, presumably due an even greater role of the host microenvironment (40, 41), Exploration of these other mechanisms of resistance is complementary to further exploration of cancer-cell genetic heterogeneity. Any systems approach taking all these mechanisms into account will be incomplete if the genetic component remains incompletely explored.

Ultrarare subclones, even if reproducibly observed across malignancies, are not universally agreed to be of clinical importance. A recent study of pancreatic cancer metastases notes general similarity among driver mutations between metastases (42). However, the sequencing was not (and could not feasibly be) performed at a depth sufficient to characterize rare subclonal variants within any one metastasis, nor within a sufficient number of metastases to determine if rare metastases have different drivers. In any case, driver mutations are likely to be more heavily selected, and thus less heterogeneous, compared to neutral mutations. In another counterargument to the importance of rare subclones, an influential approach to EGPM, adaptive therapy (23), assumes a dominant sensitive subclone can keep resistant subclones at bay due to competitive interactions, especially as a lesion grows to near its carrying capacity. However, we have pointed out that most mortality in cancer is associated with widespread microscopic metastases infiltrating organs rather than large lesions at their carrying capacity (43). Cancer is not confined to a small number of radiologically measurable lesions, and indeed exclusive or predominant reliance on radiographic assessments in large lesions may confound efforts to prevent future relapses. Microscopic lesions do not even call for angiogenesis because they are supplied with nutrients through diffusion, so competitive interactions may be limited (44). Indeed, instances where different subclones potentiate each other’s growth, rather than competing, have been observed (45). Finally, there is evidence that in some cases therapy resistance imposes a fitness cost on a cell, and in the event of competitive interactions, may lead to the extinction of the resistant subclone. In this context, we note that at the mutations rates we have measured, nearly 1000 copies of any variant will be generated with every population doubling for a cancer whose total volume across all lesions is 1 cm^3^ (1, 2), making elimination of subclones less likely, especially if the variant is repeatedly renewed. The fitness cost of a mutation is highly dependent on the genetic and microenvironmental context (46), and examples in which the resistant subclone was more fit than the original exist (47). Outgrowth of resistant subclones that were rare at diagnosis has also been documented (48). For these reasons, we believe the clinical significance of ultrarare subclones requires further investigation and may be substantial.

Our simulation of dynamic precision medicine at different limits of detection for rare subclones shows that, while DPM can be beneficial even at typical clinical sequencing sensitivities, its benefit is significantly reduced if the presence of resistant subclones below the level of detection is ignored. This is not surprising, given that we have shown that resistance to any one therapy is pre-existing (1), and that highly adaptive, proactive elimination of singly resistant subclones before they evolve further multiply resistant variants is the underlying principle of DPM.

In contrast, if the VAF-PDF is employed at diagnosis to give a probability weighted non-zero distribution below the limit of detection at diagnosis, most of the benefit of DPM is restored. The initial VAF-PDF is projected forward by the DPM mathematical model as the adaptive, personalized sequence of therapies is pursued. The method allows more prompt proactive elimination of subclones before they have time to evolve additional resistances in sub-variants. It requires treatment of subclones before detection, based on their likelihood of being present, a concept not unlike maintenance therapy. But in this case a quantitative model allows personalization and optimization of the timing and nature of maintenance therapy. The fact that the VAF-PDF outperforms other common statistical distributions for this purpose is theoretical evidence for the validity of the VAF-PDF. Experimental validation is challenging due to limitations in rare subclone detection, but work is ongoing.

Simulating the performance of DPM and other EGPM approaches in the presence of limited or incorrect input information (“mis-specification simulation”) is critical to determining the robustness of DPM and other EGPM paradigms in realistic clinical situations. We have previously performed mis-specification simulations with incorrect input information about drug sensitivities, finding that DPM benefit is surprisingly robust (49). Unlike sensitivity analysis, which examines how model outcomes change with varying parameters (50–52), mis-specification models the assumption that incorrect parameters are presumed accurate, leading to treatment decisions based on these mis-specified values. When discrepancies arise, the model user must find a way to address these differences. Furthermore, once treatments are administered, the dynamics of the cancer cell population are permanently altered by the treatment, making exact reversibility impossible. As a result, methods must be developed to recalibrate or measure the components and parameters within the model to adjust treatment accordingly (see Fig. 5 and *Materials and Methods*). Our approach offers an initial exploration of these challenges, and the insights gained could be valuable for applying model-based treatments in clinical practice.

**Figure 5.**
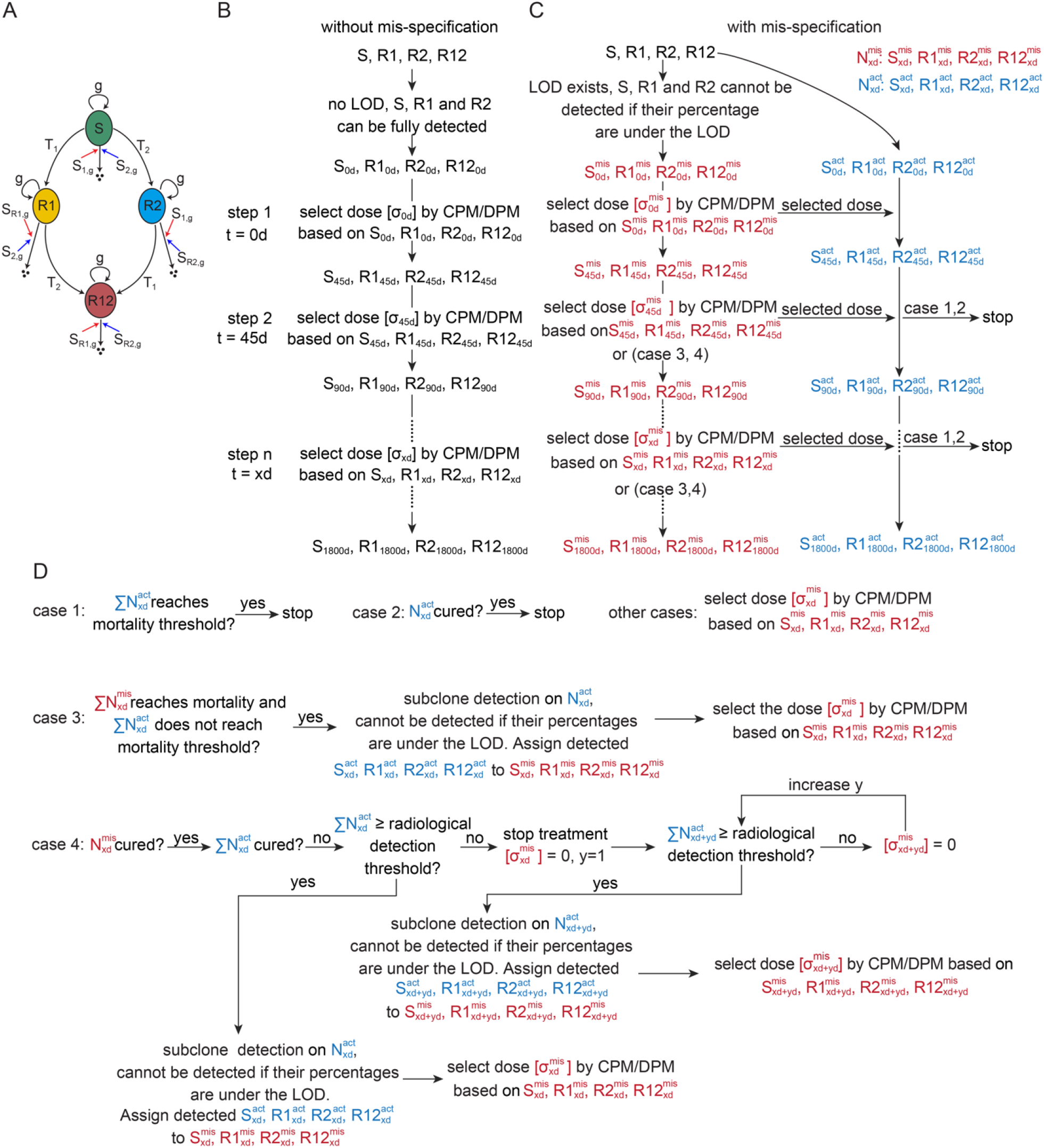
Schematic representation of the structures of the mathematical model and workflow for conducting simulations with and without mis-specification. Each simulation timestep is 45 days. The total simulation period is 1800 days, resulting in 40 steps. The number of R12 cells is 0 at t=0. LOD stands for limit of detection and d represents days. (*A*) The schematic illustrating the dynamics of different cell states and their transitions in the presence of drug 1 and drug 2, which provides the structure for the simulation. Arrows include self-replications, degradations and mutations from or to designated states. g represents the per day net growth rate; S_1,g_ represents the sensitivity of S cells to drug 1; S_2,g_, is the sensitivity of S cells to drug 2; S_R1, g_ is sensitivity of R1 cells to drug 1; S_R2, g_ is sensitivity of R2 cells to drug 2; T_1_ is the mutation rate from S to R1 cells; T_2_ is the mutation rate from S to R2 cells. (*B*) Workflow for the simulation without mis-specification. LOD does not apply, and R1 and R2 cells can are fully detectable. The drug dosage at each step *σ*_xd_] is determined by CPM or DPM. S_xd_, R1_xd_, R2_xd_ and R12_xd_ denote the cell number of S, R1, R2 and R12 at x days, respectively. (*C*) Workflow for the simulation with mis-specification. At t=0, R1 and R2 cells are undetectable if their proportions in the total cell number are below the LOD. Depending on the chosen estimation methods, R1 and/or R2 cells would be mis-specified to varying degrees. 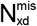 represents the mis-specified number of four cell populations at x days. 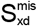, 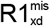, 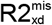 and 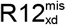 denote the mis-specified S, R1, R2 and R12 cells at x days, respectively. 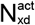 represents the actual number of four cell populations in the virtual patient at x days. 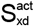, 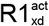, 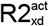 and 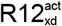 denote the actual S, R1, R2 and R12 cells in the virtual patient at x days, respectively. (*D*) Description of the cases at each timestep in the mis-specification simulation of (*C*), yd represents an additional y day of continuous simulation without treatment (detailed in *Materials and Methods*).

Genetic heterogeneity within the cancer cell population is only one aspect of the complexity of cancer. Yet, in a recent study, early intervention with the next-generational oral selective estrogen receptor degrader (SERD) camizestrant to target a minor *ESR1* mutant subclone detected by liquid biopsy resulted in significant benefit with a hazard ratio for progression-free survival of 0.44 (53). The mathematical modeling approach herein may have resulted in personalized intervention in additional patients in whom *ESR1* mutations were not detected, on a probabilistic basis based on their other evolutionary dynamic measurements at diagnosis and one or more timepoints thereafter. This may have substantially broadened the population that could have benefited from this novel SERD.

## Materials and Methods

### The probability distribution of subclonal prevalence for neutral mutations

Define *ρ* as the VAF of mutant allele(s) at a given DNA base, *ϕ*(*ρ*) represents the probability distribution of VAF. The cumulative probability of a VAF between *ρ* _*min*_ and *ρ*_*max*_ is:

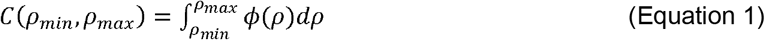

Define *N* as the total number of DNA molecules in the cancer. is the mutation rate constant, number of mutations per base per “effective” cell division (1, 2). *ρ*(*N*) is the value of *ρ* corresponding to a given value of *N*. This corresponds to the resultant value of *ρ* for a neutral mutation occurring at a particular time in the cancer evolution when the cancer burden is *N*_*cells*_ = *N*/2 cells, assuming diploid cells on average. *N*(*ρ*) is the value of *N* corresponding to a particular value of *ρ*. This corresponds to the total cancer burden when a newly occurring neutral mutation will have the VAF equal to *ρ*. Fig. 4 shows the value of *ρ* as a function of *N*_*cells*_, as previously derived (1,2).

*ρ*_*min*_ is the minimum value of for a given range of integration. For neutral mutations, *ρ* ≥*k*. Thus, when considering all possibilities *ρ*_*min*_ =*k. ρ*_*max*_ is the maximum value of *ρ* for a given range of integration. In the total tumor, *ρ*_*max*_ = 0.5 (all cells, assuming heterozygous mutations in a diploid founder cell). Under the case of undetected mutations below the limit of detection (LOD), *ρ*_*max*_ = LOD.

The earliest mutations in the tumor have higher allele fractions because the vast majority of their progeny will retain their mutations. For example, heterozygous neutral passenger mutations in a diploid founder cell will remain in nearly every cell in the tumor. At later times, with more total cells in the tumor, the allele fraction gradually approaches the limiting allele fraction *k*. The allele fraction cannot go below *k* because the DNA replication machinery sets *k* as a limiting error rate for any new base inserted. When *N*> 1/*k*, the same site will thus be mutated in multiple different daughter cells within each cell generation, on average *Nk* cells.

We took several factors into account in deriving an expression for *ϕ*(*ρ*). First, the expected fraction of genome sites that are mutated in association with the net addition of 1 new DNA molecule is proportional to *k*, the mutation rate constant. Second, in association with an infinitesimal change in VAF denoted as *d ρ*, there is a change in the total number of DNA molecules in the tumor denoted as *dN*(*ρ*). The expected fraction of genome sites mutated with the net addition of *dN*(*ρ*) DNA molecules is proportional to *dN*(*ρ*). Finally, it is important to note that at any given time a number of genome sites will already have been mutated earlier and their allele fraction will be determined by the first timepoint at which they mutated. Only previously unmutated sites are available for new mutations. Thus, the expected number of mutated sites that will be mutated at VAF *ρ* is also proportional to the number of sites still unmutated at the timepoint when the tumor has grown to *N*(*ρ*) cells. Then we have:

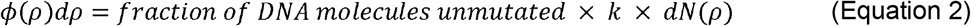

We used the fraction of genome bases unmutated when the cancer reaches a total number of DNA molecules of *N*(*ρ*) is given by the zero term of the Poisson distribution with mean *k* x *N*(*ρ*) (1, 2). The fact that this quantity obeys the Poisson distribution was verified experimentally (1, 2)

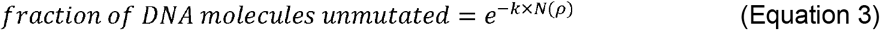

Substituting equation 3 into equation 2, and dividing both sides by *d ρ*:

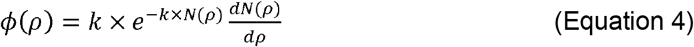

In (1, 2), we derived an expression for *ρ* (*N*) as:

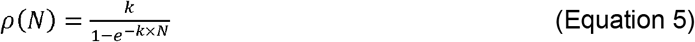

We have previously shown the limit of this expression is 1/*N* when *N* is small, or 0.5 when there is a single diploid founder cell. Similarly, it can be seen that the expression asymptotically approaches the limit *k* as the number of DNA molecules *N* becomes large, as in a radiologically measurable cancer. Solving equation 5 for *N*:

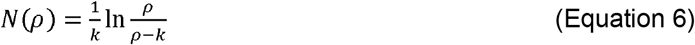

Then, multiplying both sides of equation 6 by −*k*, and raising *e* to the power of the left and right side respectively:

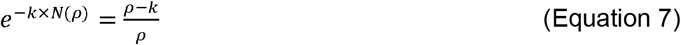

Differentiating equation 6 with respect to *ρ*:

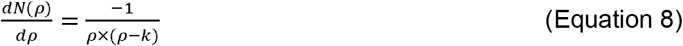

Substituting equation 7 and 8 into equation 4 gives the expression for the probability distribution of *ρ*:

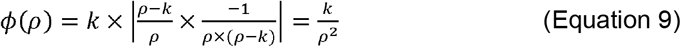

Integrating equation 9 from *ρ*_*min*_ to *ρ*_*max*_, the cumulative probability *ρ*_*cumul*_ that the value of *ρ* will be between the limits *ρ*_*min*_ and *ρ*_*max*_ :

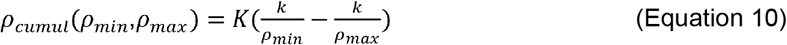

where *K* is a normalization constant to satisfy the boundary condition *ρ*_*cumul*_ =1 when *ρ*_*min*_ and *ρ*_*max*_ are at the extreme values *k* and 0.5, respectively, leading to 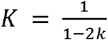, or:

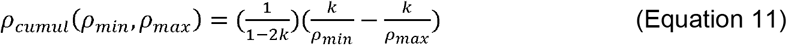

Equation 11 is used to generate Table 1 with *ρ*_*min*_ = 7.1 x 10^−7^ and *ρ*_*max*_ = 0.5.

The average VAF for neutral mutations can be calculated by summing all possible values of VAF from *k* to 0.5, weighted by the probability distribution function:

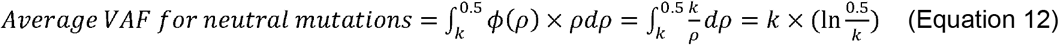

Using = 7.1 × 10^−7^ we get an average VAF of 9.4 × 10^−6^, or for a cancer burden of 10^9^ cells, an average subclonal size of 9400 cells per 10^9^ cells, as would be expected from a single 1 cm^3^ lesion.

### Mathematical model and simulation

Simulation studies are conducted to evaluate the effectiveness of two treatment methods for two drugs on two virtual patient sets, each containing 3083233 virtual patients. Each virtual patient is described by a unique set of input parameters describing their individual subclonal heterogeneity, evolutionary dynamics, and drug sensitivities at diagnosis. Virtual patient set one was constructed with the methods used in our previous work (20). There are four cell subclones in the model: S, R1, R2 and R12 cells. S cells are sensitive to both drugs, R1 cells are resistant to drug 1 and sensitive to drug 2, R2 cells are resistant to drug 2 and sensitive to drug 1, and R12 cells are resistant to both drugs. The initial total cell population is 5 × 10^9^, roughly equivalent to a 5 cm^3^ lesion. The initial cell numbers for R1 and R2 cells are set by parameters R1_ratio_ and R2_ratio,_ representing the ratio of R1 cells to the total cell number and R2 cells to the total cell number, respectively. The schematic representation of the structures of the mathematical model is shown in Fig. 5*A* (*see SI Appendix* for model details). Virtual patient set two is the same as virtual patient set one, except that R1_ratio_ and R2_ratio_ are sampled from the derived VAF-PDF equation 9 multiplied by 2 to account for the diploid nature of most of the cells, and assuming that resistance mutations are dominant. We implemented the simulation using JetStream2 (54), a resource from the Advanced Cyberinfrastructure Coordination Ecosystem: Service & Support (ACCESS) program (55). Kaplan-Meier analyses were performed using the survival analysis python library, lifelines (56).

### Treatment methods

Current Personalized Medicine (CPM): Initially, treat the virtual patient with the most effective drug based on the predominant subclone identified by molecular characterization. Define a nadir as the minimum of the total cell number during the period when the current treatment is maintained. At t = 0, nadir is equal to the initial total cell number which is 5 ×10^9^. Change the drug if either of the following events happens: (i) the total cell number reaches twice the nadir or (ii) the total cell number reemerges from a level below the detection threshold. If either (i) or (ii) occurs and another drug has not been used, switch to a different drug, meaning that each drug is used only once. Update the nadir at each timestep.

Dynamic Precision Medicine (DPM): Minimize the number of incurable cells (R12 cells) using the DPM equations (*SI Appendix)* by selecting the treatment option which projects the lowest number at the next observation timepoint, unless there is an immediate clinical risk requiring cytoreduction. In real clinical applications, the judgment of when immediate cytoreduction is necessary would be made by the clinical expert. In the simulation, we set a threshold for the number of cells in the total cancer above which cytoreduction is considered to be a priority, and if cytoreduction is a priority, the selected treatment is the one projected to minimize the total cell number at the next observation timepoint. Otherwise, at each timestep, select *σ* = (*σ*1, *σ*2), the dosages of two non-cross resistant drugs or drug combinations relative to their single agent doses, from (1, 0), (0, 1) and (0.5, 0.5) to minimize the predicted number of R12 cells, provided the total cell number does not exceed the threshold value, in this case 10^11^, an optimum determined empirically in prior work (20). Note that we assume the drug doses must be reduced in combination due to toxicity, a very common scenario. In a real application, the degree of dose reduction when both drugs are given in combination should have been determined by a Phase 1 clinical study. If not, the combination option would not be available for use in DPM. R12 cells are resistant to both drugs and are often incurable. Preventing the formation of R12 cells helps maintain the possibility for long-term survival or cure. If the total cell number exceeds 10^11^, the arbitrary threshold for prioritizing cytoreduction, select the (*σ*1, *σ*2) from (1, 0), (0, 1) and (0.5, 0.5) to minimize the predicted total cell number. DPM strategy is considered significantly better than CPM if it shows an absolute improvement of at least 56 days (8 weeks) and a relative improvement of 25% in survival time compared to the DPM strategy. This is analogous to the typical minimum improvement often regarded as clinically significant in randomized phase 3 trials in cancer (20).

### Workflow and explanation of the execution of the mis-specification simulation

As shown in Fig. 2*B* and *C*, 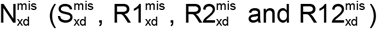 represents the number of four mis-specified cell populations at x days. 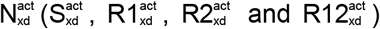 represents the actual number of four cell populations at x days. The number of R12 cells is assumed to be zero among all virtual patients at t=0, since DPM is not expected to be useful once an R12 subclone has formed. Each simulation timestep is 45 days. 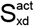, 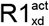, 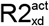 and 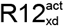 are assigned to 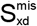, 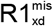, 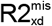 and 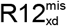, respectively, based on whether they can be detected under the limit of detection (LOD), which is a limitation on the detection of subclones. Because the cells are diploid and we hypothesize that subclones cells are heterozygous, R1 and R2 cell populations cannot be detected if their percentages in the total cell number are below 2 × LOD. The possible LOD values are 10^−6^, 10^−5^, 10^−4^, 10^−3^, 10^−2^ and 10^−1^. For example, if LOD equals 10^−2^, R1 and R2 cells cannot be detected if their percentage in the total cell numbers below 2 × 10^−2^. If the fraction of R1 cells and/or R2 cells in the total cell number is below the LOD, it is estimated using one of four methods:

1. estimation method 0: the fractions are set to 0, which means the resistant subclone is ignored.
2. estimation method max: the fractions are set to the possible maximum, which is 2 × LOD.
3. estimation method loguni: the fractions are sampled from a log-uniform distribution with PDF:

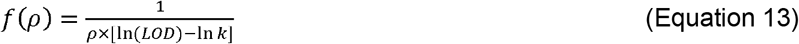
4. estimation method: VAF-PDF, the fractions are sampled from the derived PDF:

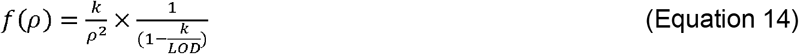

*k* = 7.1 × 10^−7^ is the mutation rate constant for both equation 4 and 5. Equation 14 is obtained by multiplying equation 9 by the normalization constant *K*, where *K*= 1/(1 – *k*/*LOD*) in equation 10 under the condition *ρ*_*min*_ = and *ρ*_*max*_ = *LOD*. The simulation setting mimics a practical drug treatment scenario. 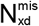 represents the measured cell populations observable by physicians, d: days, (indexed by the variable x, representing xd days). 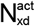 represents the actual cell populations in the patients, d: days. Using the detected or estimated cell populations 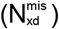, physicians decide the drug dosage based on either CPM or DPM and apply it to the patients 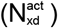. The cell populations then increase or decrease according to the drug dosage, and the simulation progresses to the next timestep. In Fig. 2*C* and *D*, except case 1 to case 4, drug dosage is determined by CPM or DPM based on 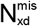 at each timestep and is applied to both 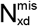 and 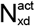. Then the simulation progresses to the next timestep. 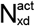 will not be re-estimated unless a specific case occurs (shown in Figs. 5*B-D*).

Case 1: Stop the simulation if the sum of 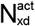 reaches the mortality threshold, resulting in actual death of the virtual patient.

Case 2: Stop the simulation if the actual virtual patient is cured, meaning each cell population in 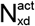 is smaller than 1. However, treatment may continue unnecessarily if the mis-specified values do not meet this criterion.

Case 3: If the sum of 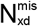 reaches mortality threshold but 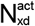 does not, meaning the virtual patient has not actually died, perform subclone detection on 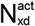, limited by the LOD, and assigned the detected subclone numbers to 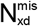. The 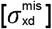 is decided based on 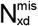 and the simulation progresses to the next timestep.

Case 4: If 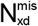 is cured but 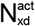 is not, and if the sum of 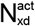 is greater than or equal to the radiological detection threshold, meaning the mis-specified estimate of cure is detectably incorrect, perform subclone detection/estimation. After subclone detection, assign the detected subclone numbers to 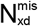, select the 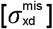 based on 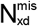, and then progress to the next timestep.

If 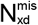 is cured but 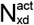 is not, and if the sum of 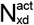 is smaller than the radiological detection threshold, stop the treatment by setting 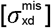 to 0. Continue the timesteps (indexed by the variable y, representing yd days) allowing the cell population to increase. Once the sum of 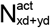 reaches the radiological detection threshold at yd days, perform subclone detection on 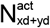 and assign the detected subclone numbers to N 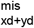, select the 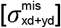 based on 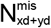 and then progress to the next timestep. Note the algorithm may be modified if a more sensitive method of detection is available based on a liquid biopsy or other technology.

## Acknowledgments

This work was funded by the Department of Defense (DoD) Breast Cancer Research Program Awards W81XWH-20-1-0759 and W81XWH-20-1-0760 (to R.B.R. and R.A.B., respectively), and by the American Cancer Society Discovery Boost Grant DBG-24-1258845-01-CDP (to R.B.R. and R.A.B.). R.A.B. was partially funded by the National Institutes of Health Cancer Center Support Grant (CCSG) 5P30CA051008. C.H.Y. was partially funded by a grant (108-2118-M-001-001-MY2) from National Science & Technology Council in the Republic of China (Taiwan). This work used Jetstream2 at Indiana University though allocation MTH240007 from the Advanced Cyberinfrastructure Coordination Ecosystem: Services & Support (ACCESS) program, which is supported by National Science Foundation grants #2138259, #2138286, #2138307, #2137603, and #2138296.

## Author Contributions

W.H. and R.A.B. designed research; W.H., and R.A.B. performed research; W.H., R.A.B. and M.D.M. analyzed data; W.H., and R.A.B. wrote the paper; M.D.M, C.H.Y., R.B.R, R.A.B. reviewed and edited; R.B.R and R.A.B provided funding acquisition, project administration and resources.

