## Supplemental file for "Probability Distribution for Rare Neutral Mutations in Cancers and Application to Dynamic Precision Medicine of Cancer"

This PDF file includes:

- Supporting text
- Figures S1 to S8
- Tables S1
- SI References

### Supporting Text

#### Additional Results for Figure 3

Fig. S3 compares several clinical metrics to assess the impact of mis-specification on clinical outcomes, accounting for increasing LODs and estimation methods. For virtual patient set one, the median survival times for the DPM strategy consistently decrease as the LOD threshold increases, as shown in Fig. S3*A*. The two sampling methods, loguni (dotted line) and VAF-PDF (dashed line), consistently outperform the methods that set the subclone percentage to boundary values (0, solid line and max, dot dash line) across all LODs. Additionally, the VAF-PDF method is superior to the loguni method at all LODs. Mis-specification has a limited impact on CPM strategy as the median survival time remains largely unchanged. The hazard values shown in Fig. S3*B* confirm the trend observed in the median survival times, with the values in DPM strategy steadily increasing as the LOD rises. Method 0 results in the worst case, followed by max and loguni, VAF-PDF provide the best improvement across all LODs. The hazard values in CPM remain largely unchanged, similar to the median survival time in CPM.

The conclusions for virtual patient set two, presented in Fig. S3*C* to *D*, align with the results from virtual patient set one. An important observation to highlight is the VAF-PDF method significantly mitigates the impact of mis-specification. The median survival time shows minimal reduction, while the hazard values exhibit only slight increase as LOD rises. It suggests that, while challenging in practice, having prior knowledge of the resistant subclone percentage distribution allows for sampling potential percentages to approximate the unknown actual values, effectively compensating for experimental limitations. Fig. S3*E* and *F* display bar plots illustrating the changes in number of cases, with DPM significantly better than CPM across various estimation methods for virtual patient set one and set two, respectively. Significantly better is defined as providing at least a 25% relative improvement and at least a 2 month absolute improvement in survival relative to CPM. These findings are consistent with the clinical metrics shown in Figs. S3*A* to S3*D*, which demonstrate that increasing levels of misspecification progressively reduce the advantage of the DPM strategy over CPM as the LOD increases. Furthermore, the VAF-PDF estimation method alleviates this impact, particularly when the resistant subclone percentage distribution aligns with the VAF-PDF.

#### Virtual patient examples demonstrating how mis-specification diminishing the benefit of DPM in the presence of LOD

Figs. S4*A* to S4*F* use a virtual patient from set one to illustrate that the DPM strategy can extend the virtual patient's survival to 5 years, but this benefit diminishes in the presence of LOD. Fig. S4*A* shows the cell number dynamics under the CPM strategy without mis-specification, indicating that resistant subclones can be detected regardless of their rarity within the heterogeneous tumor population. The CPM strategy targets the dominant subclone, represented by the S cells (green line) in this virtual patient. As a result, the most effective drug, drug 1 (red), is used at t = 0. This leads to a rapid decrease in the number of S cells, eliminated by drug 1. Consequently, the total cell number quickly falls below the radiologic detection limit, indicating that the tumor becomes undetectable using standard imaging methods. However, R1 cells (yellow line), which are resistant to drug 1, continue to grow undetected by imaging, giving rise to R12 cells (maroon line) through mutation. Relapse happens at 225 days after initiating treatment with drug 1, indicating that the tumor burden has reappeared above the limit of radiologic detection. This indicates the failure of drug 1 in the CPM strategy and a switch to drug 2. At this stage, the heterogeneous tumor population consists of R1 and R12 subclones. Although drug 2 reduces the R1 cells and again brings the total tumor burden below the limit of radiologic detection, it fails to control the R12 cells, which continue to proliferate undetected. Inevitably, relapse happens again around 540 days, with the dominant subclone becoming R12. As no available drug can target this double resistant subclone, mortality occurs at approximately 720 days, or about two years. This scenario represents a potential outcome for a patient undergoing current personalized medicine, where two relapses occur and the second relapse is dominated by the emergence of a subclone resistant to one or more of the drugs used.

Fig. S4*B* shows the cell number dynamics under the DPM strategy without mis-specification. Unlike the CPM approach, which targets the dominant subclone in the cell population, DPM focuses on minimizing the R12 cells. DPM selected drug 2 over drug 1 due to the detectable presence of R1 cells, which are more sensitive to drug 2 and are only mutational step away from evolving to R12 cells that are incurable in the proposed scenario. Using drug 2 gradually reduced S and R1 cells, but R2 cells start to gradually increase. Around day 225, the number of R2 cells exceeded R1 cells, prompting the selection of a combination of drug 1 and 2. The drug combination, interleaved with a full dosage of drug 1, effectively suppressed both the R1 and R2 cells in a staggered manner and prevented the emergence of R12 cells.

Figs. S4*C* and S4*D* show the cell number changes under the DPM strategy for the same virtual patient with a LOD of $10^{-2}$ and an estimation method 0. Fig. S4*C* illustrates the mis-specified cell number changes and Fig. S4*D* presents the actual cell number changes. Drug selection decisions are based on the mis-specified cell numbers. Subclone detection at t = 0 failed to identify R1 cells because their number was below the LOD, shown in Fig. S4*D*, represented by a dotted line. In Fig. S4*C*, the tumor cells were mis-specified to be homogeneous S cells, despite being heterogeneous. Although the DPM strategy is still applied to minimize R12 cells, drug 1 is initially selected as it is more effective against S cells. However, the mis-specification causes the DPM to revert to a CPM-like approach during this critical phase of initial drug selection. After three timesteps of drug 1 treatment, the virtual patient was incorrectly assumed to be cured, as all the mis-specified cell numbers had fallen below 1, leading to the cessation of drug treatment. However, R1 cells continued to proliferate unnoticed and rapidly emerged shortly after the drug treatment was discontinued. A subclone detection performed at that timepoint revealed that R1 cells had become the dominant subclone in the tumor. Although treatment the drug 2 was then started, the initial drug was mis-selected and the critical opportunity to prevent the emergence of R12 cells was missed, as by that point the R1 cells had evolved a sub-population of R12 cells. As a result, the survival time for this virtual patient was significantly reduced to less than 900 days. The cell number dynamics under the CPM strategy with an LOD of $10^{-2}$ and mis-specification method 0 are shown in Figs. S5*A* and S5*B*. The survival time for this virtual patient under CPM with mis-specification remains approximately 720 days, as the initial drug selections were not affected by the mis-specification. Fig. S4*E* shows the total cell number changes for CPM (red line) and DPM (solid blue line) without mis-specification, and the actual total cell number changes of DPM (dashed blue line) under mis-specification. CPM initially results in a shaper decrease in cell numbers compared to DPM. But two relapses occur due to the emergence of drug resistant subclones and ultimately leading to mortality. The mis-specifications of subclones diminishes the advantages of DPM, resulting in a survival time similar to that of CPM. As shown in Fig. S4*F,* R12 cells cause the virtual patient's mortality under both the CPM strategy without mis-specification and the DPM strategy with mis-specification.

Figs. S4*G* to S4*L* present another case from virtual patient set two to show that while the DPM strategy can rapidly achieve an apparent cure, relapse can occur over a year after treatment cessation due to the presence of the LOD. Fig. S4*G*, similar to Fig. S4*A*, shows that CPM initially achieves a sharper decrease in cell numbers, but two relapses occur due to the emergence of drug resistant subclones ultimately resulting in mortality. In Fig. S4*H*, we show the cell number dynamics under the DPM strategy without mis-specification. Unlike the CPM approach, which targets the dominant subclone, DPM focuses on minimizing the R12 population. Therefore, DPM selects drug 2 initially to prevent the emergence of R12 cells, since choosing drug 1, as seen in the CPM case, would promote the formation of R12 cells. In the first step, selecting drug 2 leads to an increase in R2 cells at the first step. Then in the second step, DPM selects a drug combination that reduces R2 cells while eliminating R1 cells. In the third step, DPM selects drug 1 to fully eliminate the remaining R2 cells, rapidly achieving a cure for this virtual patient within only three timesteps. In the presence of LOD, neither R1 nor R2 cells are detectable, leading to the assumption that the virtual patient was cured around three timesteps using the DPM strategy. However, the undetectable resistant subclones gradually proliferate during the drug withdrawal period. Approximately one year after the patient is presumed cured, the tumor burden reemerges and an R12 subclone develops during the drug withdrawal period. The virtual patient becomes untreatable with the available drugs and ultimately resulting in mortality over three years later. Figs. S4*K* and S4*L* illustrate the changes in total cell numbers and R12 cell numbers, respectively. The two relapses observed in the CPM and DPM strategies with mis-specification were driven by the emergence of resistant subclones, with the R12 cells ultimately causing mortality. The cell number dynamics of this virtual patient under the CPM strategy with mis-specification are shown in Figs. S5*C* and S5*D*. The survival time is reduced somewhat due to a treatment withdrawal caused by the mistakenly assumed cure.

**Mis-specification disrupting the drug scheduling in DPM**

A key principle of DPM is its proactive approach, instead of waiting for a relapse to occur, DPM introduces the second line drug (drug 2 in our model) earlier to reduce the risk of future relapse. To evaluate how drug 2 is utilized under CPM and CPM, both with and without mis-specification, we calculated three metrics for the two virtual patient sets. Figs. S6*A* and 6*C* present the average timestep for introducing drug 2 in virtual patient sets one and two, respectively. In both sets, DPM initiates drug 2 earlier on average compared to CPM when no mis-specification is present. In virtual patient set two, CPM introduces drug 2 even later than in set one, as the initial population of resistant subclones is lower, having been sampled from the VAF-PDF rather than set to higher possible values as was done in set one (see Table S1). DPM delayed the usage of drug 2 nearly for all estimation methods as the LOD increased, especially when we ignore the possible existence of rare resistant subclones by using estimation method 0 (blue solid line). Consistent with previous results, the VAF-PDF estimation method (blue dashed line) alleviates this impact with a nearly unchanged average drug 2 introduction timestep across the LODs. The average drug 2 introduction timestep is moved forward for CPM with estimation method max (red dash dot line) in virtual patient set one, possibly because setting the maximum value of the resistant subclone in the mis-specified cell population results in use of drug 2 earlier than is optimal.

Figs. S6*B* and 6*E* present the average drug 2 intensity for virtual patient sets one and two, respectively. In both virtual patient sets, DPM used a higher average dosage of drug 2 compared to CPM when no mis-specification was introduced. Across both sets, all estimation methods resulted in a reduction of the drug 2 dosage in DPM. Estimation method 0 led to a sharp decrease in DPM drug 2 usage while method max gradually restored the dosage to levels comparable to the no mis-specification case as the LOD increased, by assigning the resistant subclone to its maximum possible proportion. For CPM, both the max and loguni estimation methods led to an increase in the average drug 2 dosage for both virtual patient sets as the LOD increased. This is because assigning a higher proportion to resistant subclones extended the survival time for some virtual patients under CPM, contributing to a higher average drug dosage. The drug 2 time weighted average dose intensity shown in Figs. S6*C* and 6*F*, reflects how late drug 2 is administered, incorporating both the timing of its introduction and its dosage intensity. Without mis-specification, this value is similar for DPM compared to CPM in virtual patient set one, indicating continued use of drug 2 throughout the patient's course for DPM. However, in set two, the metric is 40% higher for CPM as CPM introduces drug 2 much later in set two, where the patient cohort begins with lower values of R1 cells since the VAF-PDF is concentrated at low values. Estimation method 0 leads to an increase in this metric for DPM in both parameter sets, indicating delayed drug 2 usage. In contrast, estimation method max results in a decrease for CPM in both virtual patient sets, indicating earlier drug 2 usage.

The percentage of identical DPM dosages with and without mis-specification for virtual patient sets one and two are shown in Figs. S7*C* and 7*D*, respectively. As expected, these percentages decrease as the LOD increases. Among the estimation methods, loguni and VAF-PDF generally maintain the highest consistency in drug selection between the mis-specified and non-mis-specified cases, particular during the initial timesteps. For comparison, the percentage of identical CPM dosages with and without mis-specification for virtual patient sets one and two are shown in Fig. S8*C* and *D*, respectively. Since CPM focuses on targeting the dominant subclone at the initial timestep, and this dominant subclone is not influenced by the LOD, the drug selection at the first timestep remains unchanged. These results shown in Fig. S6 and Fig. S7 demonstrate that mis-specification disrupts drug scheduling in DPM, thereby reducing its advantage over CPM.

**Model Parameters**

There are a total nine parameters in the model, which are:

(*i*). g represents the per day net growth rate. (*ii*). $S_{1,g}$ represents the sensitivity of S cells to drug 1 relative to the net growth rate g. That is, a full dose of drug 1 will decrease the net growth rate of S cells by the amount $S_{1,g}$ × g (*iii*) $S_{2,1}$ represent the sensitivity of S cells to drug 2 relative to drug 1. We assumed that drug 1 is more effective than drug 2 against S cells, mimicking the clinical use of drug 1 as the first-line treatment and drug 2 as the second-line treatment. $S_{2,g}$, is the sensitivity of S cells to drug 2 relative to the net growth rate g, and is calculated as $S_{1,g}$ × $S_{2,1}$. (*iv*) $R1_{1,S}$ is the sensitivity of R1 cells to drug 1 relative to the sensitivity of S cells to drug 1. $R1_{1,g}$ is sensitivity of R1 cells to drug 1 relative to the net growth rate g, and is calculated as $S_{1,g}$ × $R1_{1,S.}$ R1 cells have the same sensitivity to drug 2 as S cells. (*v*) $R2_{2,S}$ is the sensitivity of R2 cells to drug 2 relative to the sensitivity of S cells to drug 2. $R2_{2,g}$ is sensitivity of R2 cells to drug 2 relative to the net growth rate g, and is calculated as $S_{2,g}$ × $R2_{2,S.}$ R2 cells have the same sensitivity to drug 1 as S cells. (*vi*) $T_1$ is the mutation rate from S to R1 cells. (*vii*) $T_2$ is the mutation rate from S to R2 cells. (*viii*) $R1_{ratio}$ is the ratio of the initial R1 cell number to the total cell number. (*ix*) $R2_{ratio}$ is the ratio of the initial R2 cell number to the total cell number. The value ranges of these nine parameters and the discrete values sampled across the range, as shown in Table S1, are the same as those in our previous work (1). The ranges of growth rates are approximations from progression and/or survival times from indolent or aggressive cancers, while the ranges of transition rates are estimated from the most accurate and least accurate DNA polymerases, assuming from 1-100 mutational targets that might lead to a specific resistance phenotype. These ranges are meant to be approximate and inclusive (1).

**Mathematical Model**

The four different cell subclones are represented by x = (S, R1, R2, R12). All cells have the same growth rate g. The cell death rates are 0 without treatment, and the drugs work by increasing death rate. The model is expressed by a differential equation:

$$\frac{dx}{dt} = [(I + T) - diag(S_a \times \sigma)] \times g \times U(x - 1) \times x \qquad \text{(Equation S1)}$$

where $I$ is an identity matrix and $diag(\cdot)$ is an operator that places components on the diagonal entries of a zero matrix. $U(x - 1) \times x$ is the Heaviside step function that sets component values to 0 if they are less than 1. Specifically, $U(x_i - 1) = 0$ if $x_i < 1$ and $U(x_i - 1) = 1$ if $x_i \geq 1$, where $x_i$ is the component in $x, x_i \in \{S, R1, R2, R12\}$. This term ensures that fractional cell numbers (i.e., cell numbers less than 1) will not contribute to cell division. $T$ is a transition rate matrix corresponding to the mutations from all other cell types, which is denoted as follows:

$$T = \begin{pmatrix} 0 & 0 & 0 & 0 \\ T_1 & 0 & 0 & 0 \\ T_2 & 0 & 0 & 0 \\ 0 & T_2 & T_1 & 0 \end{pmatrix} \qquad \text{(Equation S2)}$$

$T(i, j)$ specifics the mutation rate from subclone $j$ to $i$. We assumed that (*i*) mutations from resistant to sensitive subclones are negligible, (*ii*) the mutation of acquiring the resistance to one drug is independent of the resistance state to another drug, and (*iii*) mutations of acquiring double resistance in one step are negligible. $S_a$ is a matrix representing sensitivities relative to the net growth rate, which is denoted as follows:

$$S_a = \begin{pmatrix} S_{1,g} & S_{2,g} \\ S_{R1,g} & S_{2,g} \\ S_{1,g} & S_{R2,g} \\ S_{R1,g} & S_{R2,g} \end{pmatrix} = \begin{pmatrix} S_{1,g} & S_{2,1} \times S_{1,g} \\ R_{1,S} \times S_{1,g} & S_{2,1} \times S_{1,g} \\ S_{1,g} & R_{2,S} \times S_{2,1} \times S_{1,g} \\ R_{1,S} \times S_{1,g} & R_{2,S} \times S_{2,1} \times S_{1,g} \end{pmatrix} \quad \text{(Equation S3)}$$

The schematic representation of the structures of the mathematical model is shown in Fig. 5*A*.

**Calculation of metrics**

We defined three drug usage metrics to evaluate the usage of drug 2:

(*i*) The average drug 2 introduction timestep is defined as $\frac{1}{N} \times \sum_{i=1}^{N} I_{drug2}$, which represents the average timestep where drug 2 was first used. $I_{drug2}$ denotes the index of the timestep where drug 2 was first used, $N$ is the total number of virtual patients and $i$ is the index of the virtual patients. Since the total number of timesteps is 40, the possible values for $I_{drug2}$ range from 1 to 40. If drug 2 has never been used for a virtual patient, $I_{drug2}$ is set to the maximum number of timesteps, which is 40.

(*ii*) The average drug 2 intensity is defined as $\frac{1}{N} \times \left(\sum_{i=1}^{N} (\frac{1}{M_i} \sum_{j=1}^{M_i} \sigma 2_j)\right)$, which represents the averaged mean drug 2 dosage intensity across all virtual patients. $\sigma 2_j$ is the drug 2 dosage at the $j$th timestep for the $i$th virtual patient, with the possible value 0, 0.5, and 1. $M_i$ is the total number of timesteps for the $i$th virtual patient, ranging from 1 to 40, and $N$ is the total number of virtual patients.

(*iii*) The drug 2 time weighted average intensity is defined as $\frac{1}{N} \times \left(\sum_{i=1}^{N} \left(\frac{\sum_{j=1}^{M_i} j \times \sigma 2_i}{\sum_{j=1}^{M_i} \sigma 2_i}\right)\right)$, representing the average time weighted mean drug 2 dosage intensity across all virtual patients. $\sigma 2_j$ is the drug 2 dosage at the $j$th timestep for the $i$th virtual patient. $M_i$ is the total number of timesteps for the $i$th virtual patient, ranging from 1 to 40, and $N$ is the total number of virtual patients.

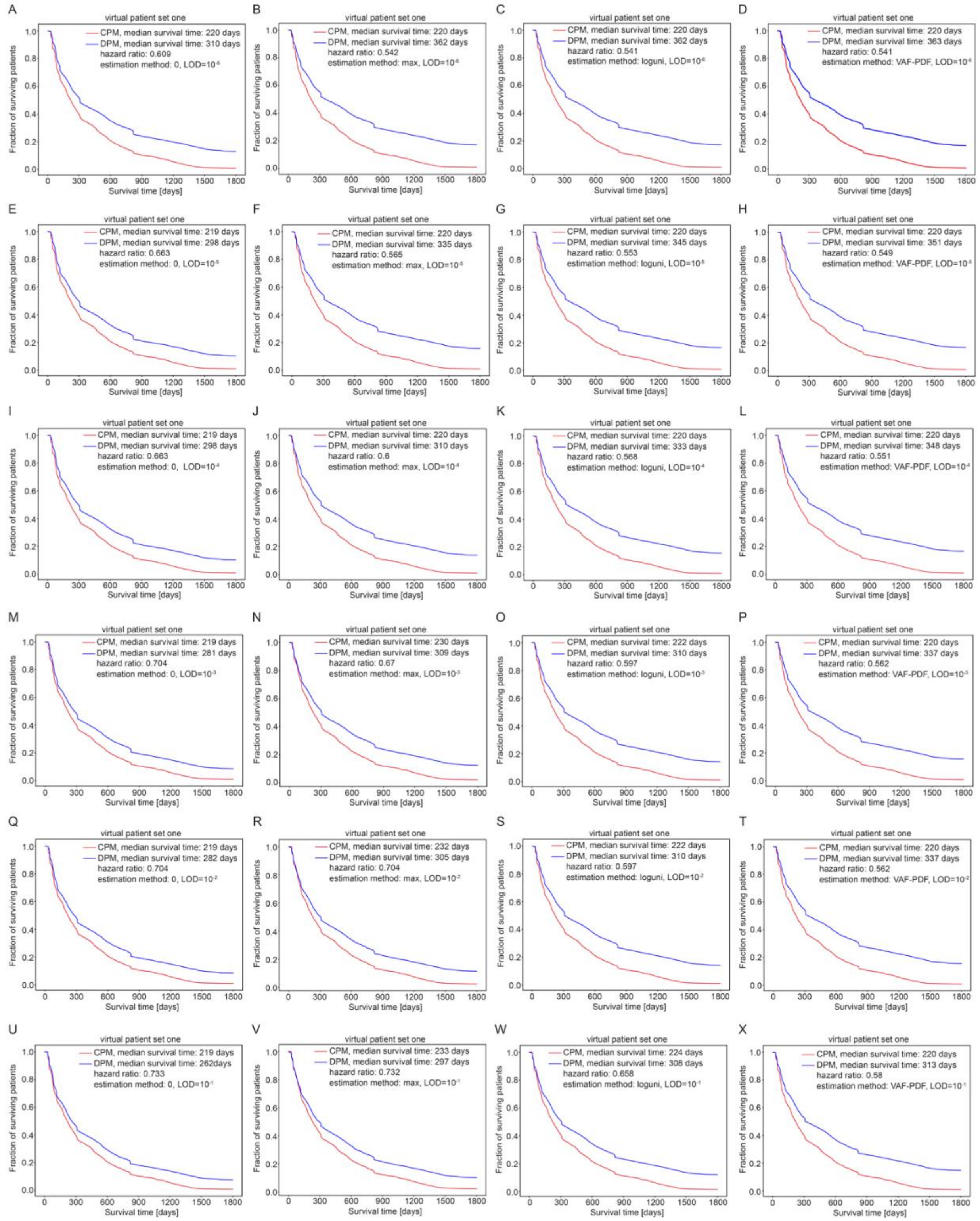


**Fig. S1.** Kaplan-Meier survival curves for CPM and DPM strategies in virtual patient set one at varying subclonal limits of detection (LOD). Virtual patient set one is the patient set used in the original DPM simulation, and varies the prevalence of the singly resistant subclones among equally spaced alternatives in a logarithmically transformed space, over a broad range (1). 3083233 virtual patients were treated with these two strategies. The x axis shows time and the y axis shows the surviving patient fraction. CPM is shown in red and DPM is shown in blue. (*A*-*D*) LOD equals $10^{-6}$. (*A*) The fractions of R1 and R2 cells below the LOD are set to 0. (*B*) The fractions of R1 and R2 cells below the LOD are set to the maximum possible value if they are undetected, i.e. at the LOD. (*C*) The fractions of R1 and R2 cells are sampled from a log-uniform distribution. (*D*) The fractions of R1 and R2 cells are sampled from the VAF-PDF (Equation 14, main manuscript, *Materials and Methods*). (*E*-*H*) Same as (*A*-*D*), except LOD equals $10^{-5}$. (*I*-*L*) Same as (*A*-*D*), except LOD equals $10^{-4}$. (*M*-*P*) Same as (*A*-*D*), except LOD equals $10^{-3}$. (*Q*-*T*) Same as (*A*-*D*), except LOD equals $10^{-2}$. (*U*-*X*) same as (*A*-*D*), except LOD equals $10^{-1}$.

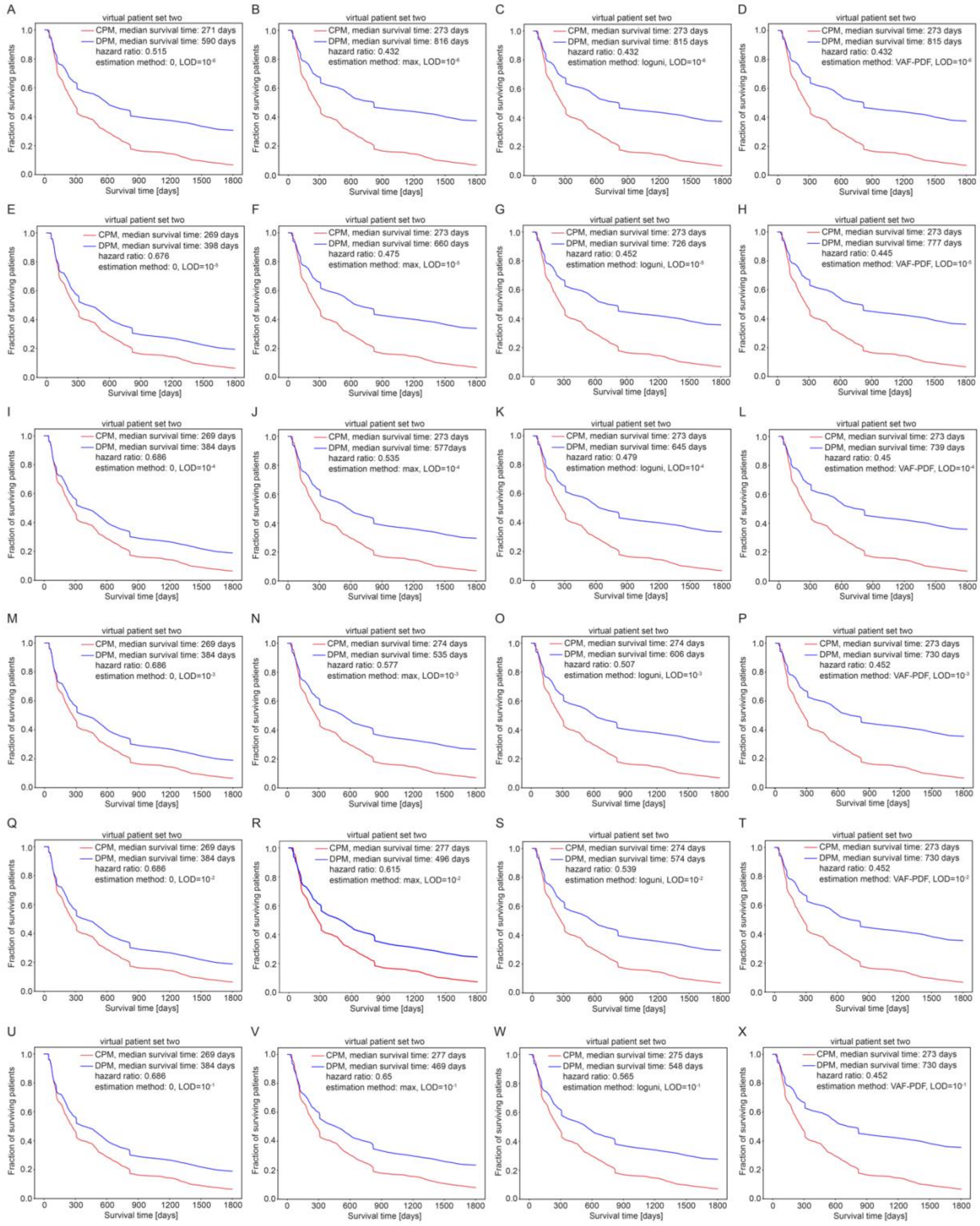


**Fig. S2.** Kaplan-Meier survival curves for CPM and DPM strategies at varying subclonal limits of detection (LOD), as in Figure S1 except using virtual patient set two. For virtual patient set two, we sample the VAF-PDF (multiplied by 2) to determine the prevalence of the singly resistant subclones. 3083233 virtual patients were treated with these two strategies. The x axis shows time and the y axis shows the surviving patient fraction.

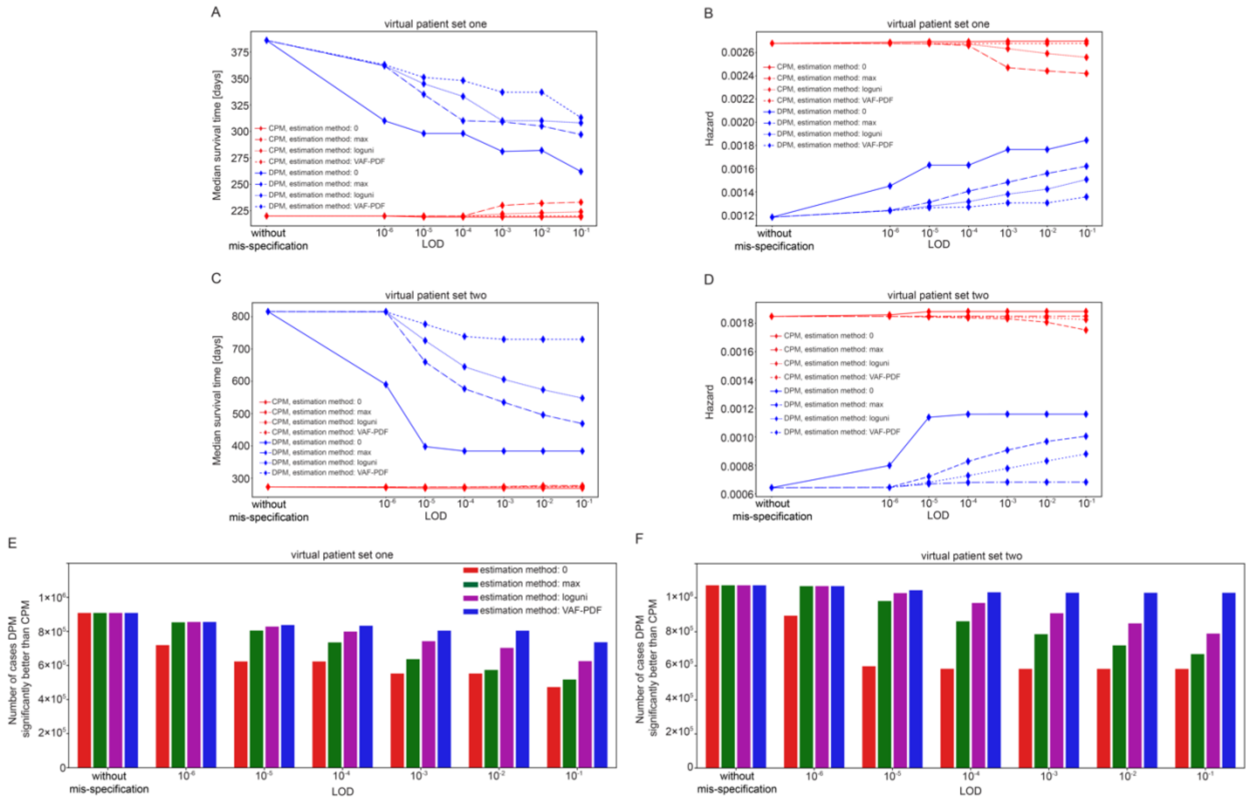


**Fig. S3.** Comparison of various metrics under scenarios with and without mis-specifications and varying LODs for virtual patient set one and two. Virtual patient set one is the patient set used in the original DPM simulation and varies the prevalence of the singly resistant subclones among equally spaced alternatives in a logarithmically transformed space, over a broad range (1). For virtual patient set two, we sample the VAF-PDF (multiplied by 2) to determine the prevalence of the singly resistant subclones. (*A*) Changes in median survival time for CPM and DPM across various estimation methods, with increasing LODs, starting from the scenario without mis-specification for virtual patient set one. (*B*) Changes in hazard values for CPM and DPM across various estimation methods, with increasing LODs, starting from the scenario without mis-specification for virtual patient set one. (*C*) Same as (*A*) but for virtual patient set two. (*D*) Same as (*B*) but for virtual patient set two. (*E*) Bar plot of changes in number of cases DPM significantly better than CPM across various estimation methods, with increasing LODs, starting from the scenario without mis-specification for virtual patient set one. DPM is scored as significantly better if it provides both a minimum 25% increase and minimum 2 month absolute improvement in median survival. Of note, DPM was never significantly inferior to CPM over these large virtual patient sets encompassing very wide ranges of all the input parameters. (*F*) Same as (*E*) but for virtual patient set two.

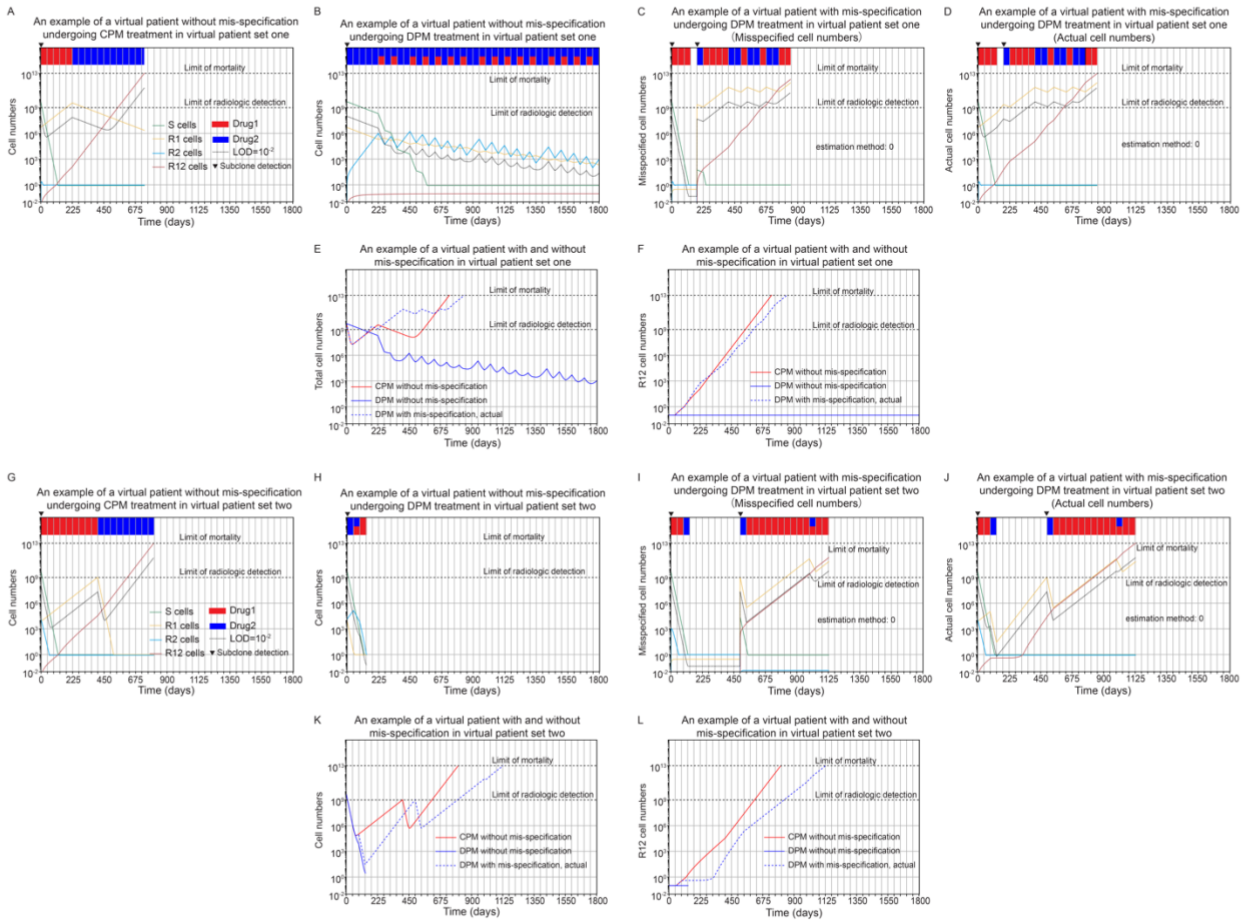


**Fig. S4.** Population dynamics of two illustrative virtual patient examples demonstrate how mis-specification diminishes the benefit of DPM compared to CPM in virtual patient sets one and two, with a LOD of $10^{-2}$ and an estimation method 0, equivalent to current practice but the least effective of the estimations simulated. Virtual patient set one is the patient set used in the original DPM simulation and varies the prevalence of the singly resistant subclones among equally spaced alternatives in a logarithmically transformed space, over a broad range (1). For virtual patient set two, we sample the VAF-PDF (multiplied by 2) to determine the prevalence of the singly resistant subclones. (*A* – *F*) A virtual patient example from set one shows that the survival time advantage of the DPM strategy, extending up to 5 years, is diminished in the presence of the LOD. (*A*) Cell number dynamics under the CPM strategy without mis-specification. The drug used at each timestep are indicated on the top of the plot, drug 1 shown in red and drug 2 shown in blue. (*B*) Cell number dynamics under the DPM strategy without mis-specification. (*C*) The misspecified cell number dynamics under the DPM strategy in the presence of mis-specification. (*D*) The actual cell number dynamics under the DPM strategy in the presence of mis-specification. (*E*) The total cell number dynamics for the CPM and DPM strategies without mis-specification, alongside the actual total cell number dynamics for the DPM strategy with mis-specification. (*F*) The R12 cell number dynamics for the CPM and DPM strategies without mis-specification, alongside the actual R12 cell number dynamics for the DPM strategy with mis-specification. The arrow at the top of the figure indicates when subclone detection occurs, either at time zero or during the treatment phase misspecification scenarios (case 4 in Fig. 5*D*). (*G* - *L*) A virtual patient example from set two shows how the DPM strategy's curative advantage is diminished in the presence of the LOD. (*G*) Same as (*A*) but for a virtual patient example from set two. (*H*) Same as (*B*) but for a virtual patient example from set two. (*I*) Same as (*C*) but for a virtual patient example from set two. (*J*) Same as (*D*) but for a virtual patient example from set two. (*K*) Same as (*E*) but for a virtual patient example from set two. (*L*) Same as (*F*) but for a virtual patient example from set two.

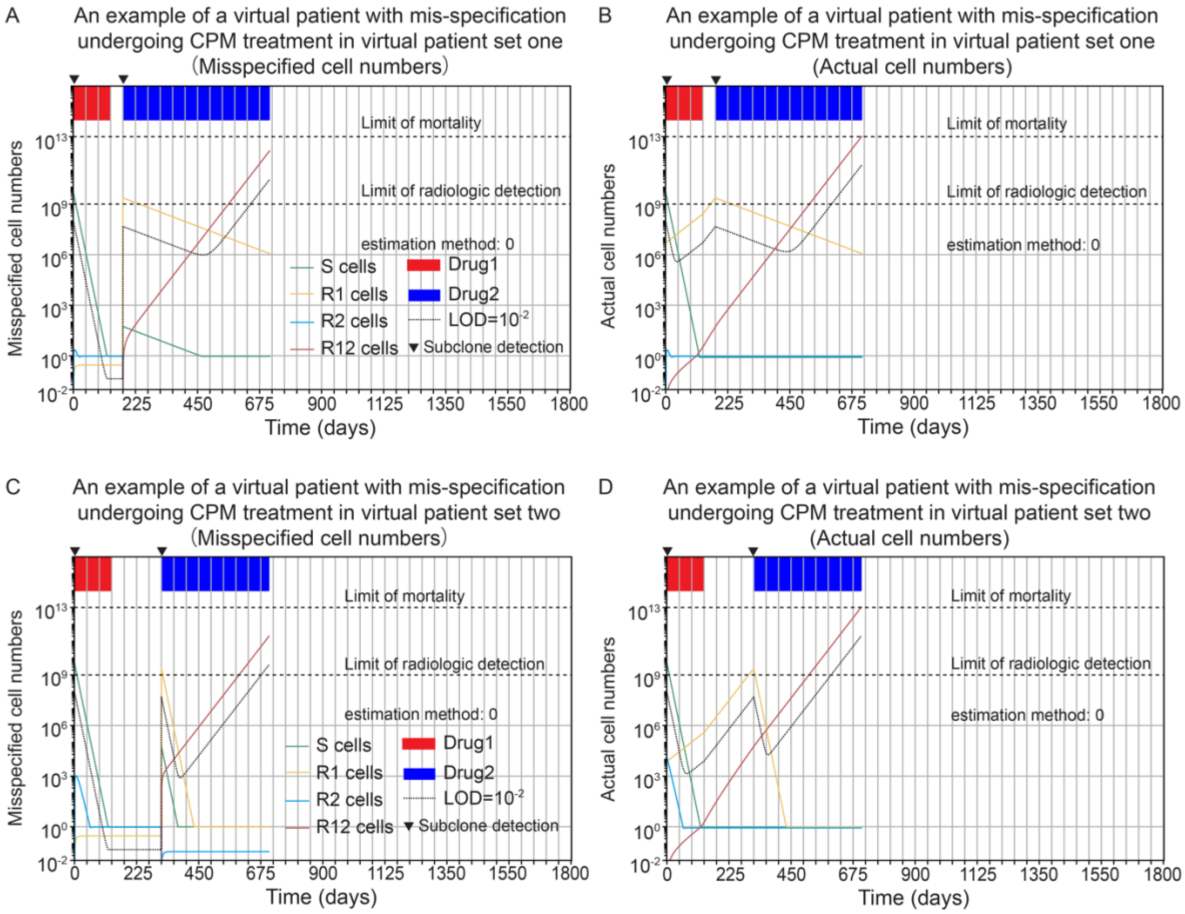


**Fig. S5.** The cell number dynamics of the virtual patient examples shown in Fig. S4, simulated under the CPM strategy with a LOD of $10^{-2}$ and using estimation method 0, in that treatment is paused between the use of drug 1 and that of drug 2 until relapse is detected. In essence, the inability to detect rare cells results in the misclassification of the patient as one without minimal residual disease. Virtual patient set one is the patient set used in the original DPM simulation and varies the prevalence of the singly resistant subclones among equally spaced alternatives in a logarithmically transformed space, over a broad range (1). For virtual patient set two, we sample the VAF-PDF (multiplied by 2) to determine the prevalence of the singly resistant subclones. (*A*-*B*) The example from virtual patient set one demonstrates that mis-specification can result in an inappropriate termination of treatment (the same virtual patient set as in Figs. S4*A*–S6*F*). (*C*-*D*) The example from virtual patient set two similarly shows that mis-specification can result in an inappropriate termination of treatment (the same virtual patient set as in Figs. S4*G*–S6*L*).

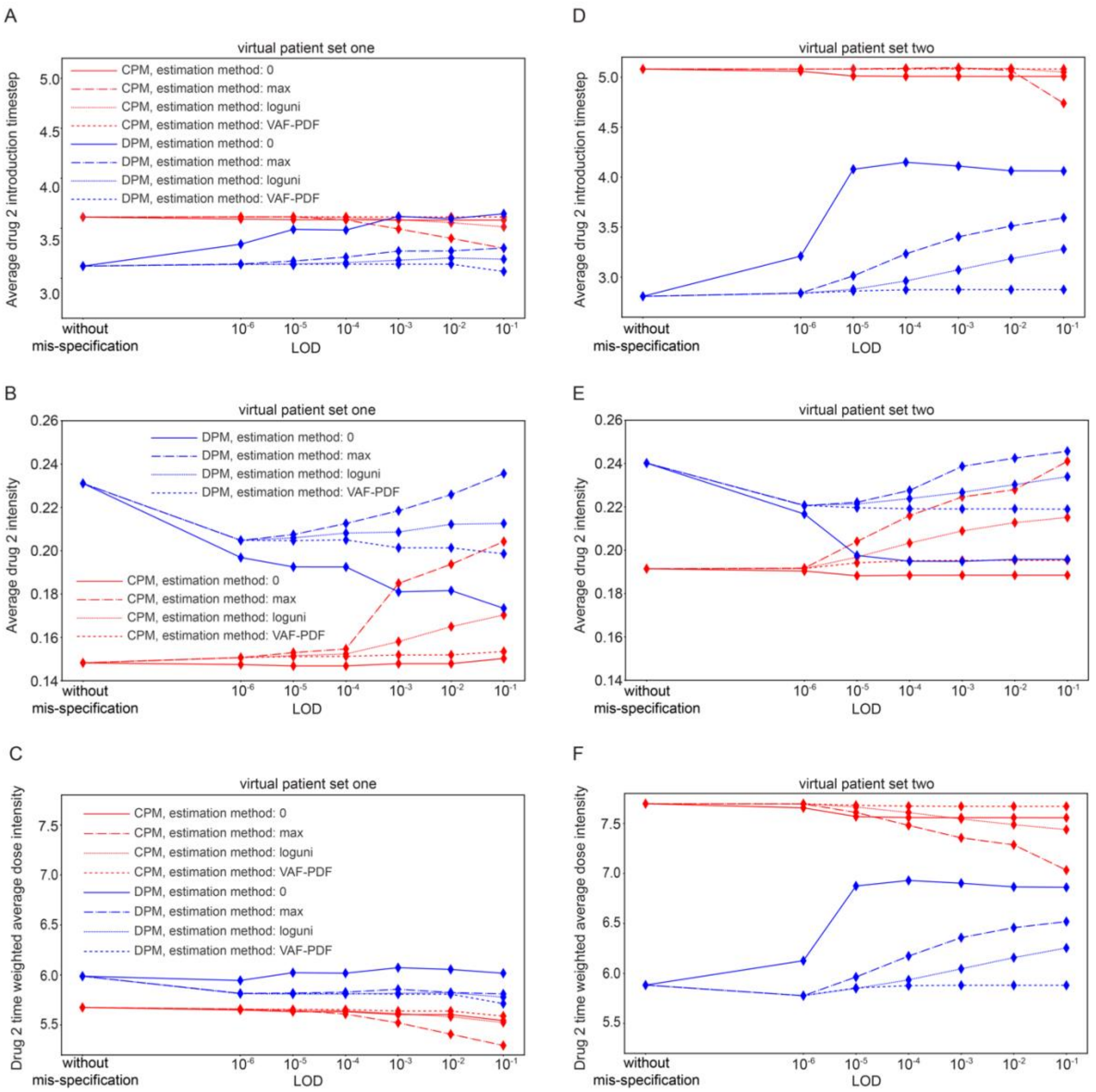


**Fig. S6.** Comparison of metrics evaluating the drug 2 use across scenarios with and without mis-specifications, shown for increasing LODs, starting from the scenario without mis-specification for virtual patient sets one and two. Virtual patient set one is the patient set used in the original DPM simulation and varies the prevalence of the singly resistant subclones among equally spaced alternatives in a logarithmically transformed space, over a broad range (1). For virtual patient set two, we sample the VAF-PDF (multiplied by 2) to determine the prevalence of the singly resistant subclones. (*A*) Changes in average drug 2 induction timestep for CPM and DPM across various estimation methods for virtual patient set one. (*B*) Changes in average drug 2 intensity for CPM and DPM across various estimation methods for virtual patient set one. (*C*) Changes in drug 2 time weighted average dose intensity for CPM and DPM across various estimation methods for virtual patient set one. (*D*) Same as (*A*) but for virtual patient set two. (*E*) Same as (*B*) but for virtual patient set two. (*F*) Same as (*C*) but for virtual patient set two.

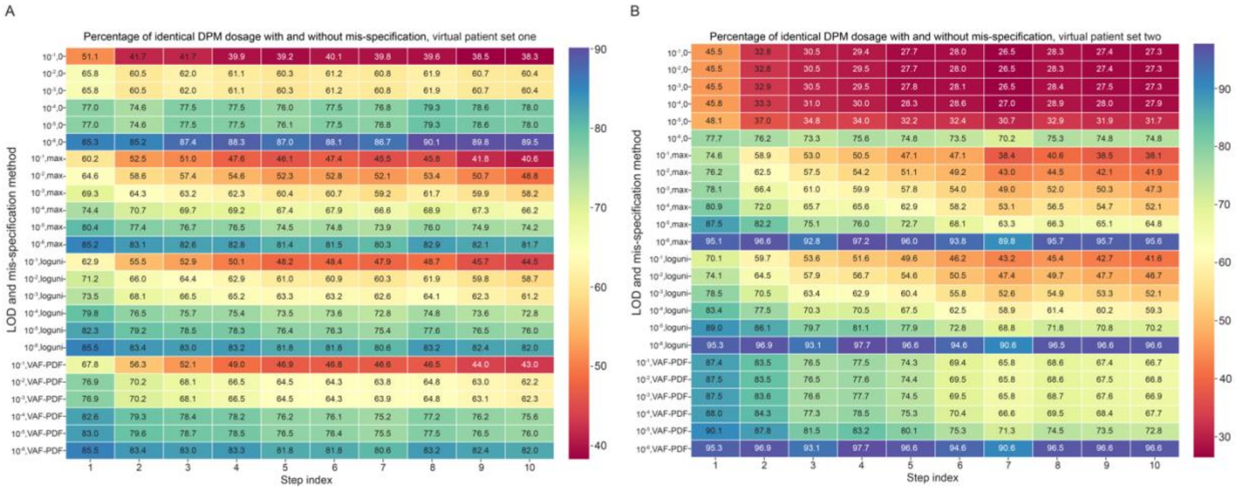


**Fig. S7.** Illustration of the drug dosage changes for DPM across scenarios with and without mis-specifications and varying LODs for virtual patient sets one and two. Virtual patient set one is the patient set used in the original DPM simulation and varies the prevalence of the singly resistant subclones among equally spaced alternatives in a logarithmically transformed space, over a broad range (1). For virtual patient set two, we sample the VAF-PDF (multiplied by 2) to determine the prevalence of the singly resistant subclones. (*A*) Percentage of identical DPM dosage with and without mis-specification across various estimation methods and LODs for virtual patient set one. (*B*) Same as (*A*) but for virtual patient set two.

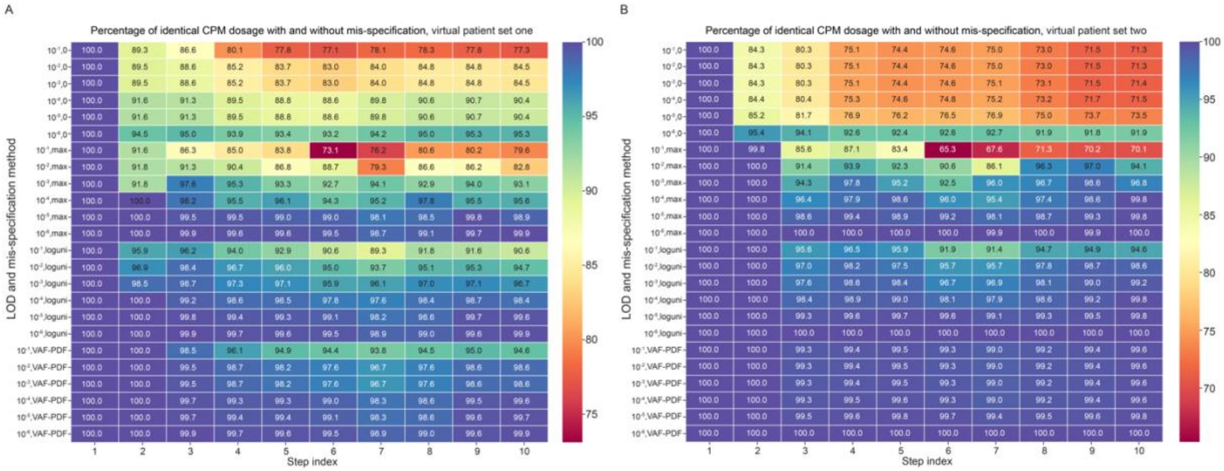

**Fig. S8.** Illustration of the drug dosage changes for CPM across scenarios with and without mis-specifications and varying LODs for virtual patient sets one and two. Virtual patient set one is the patient set used in the original DPM simulation and varies the prevalence of the singly resistant subclones among equally spaced alternatives in a logarithmically transformed space, over a broad range (1). For virtual patient set two, we sample the VAF-PDF (multiplied by 2) to determine the prevalence of the singly resistant subclones.(*A*) Percentage of identical CPM dosage with and without mis-misspecification across various mis-specification methods and LODs for virtual patient set one. (*B*) Same as (*A*) but for virtual patient set two. Virtual patient set one is the patient set used in the original DPM simulation (1). For virtual patient set two, we sample the VAF-PDF (multiplied by 2) to determine the prevalence of the singly resistant subclones.

**Table S1.** Model parameters and values

| Parameter | Description | Values |
|---|---|---|
| (*i*) g | net growth rate | 0.001, 0.002642, 0.00698, 0.018439, 0.048714, 0.128696, 0.34 |
| (*ii*) $S_{1,g}$ | sensitivity of S cells to drug 1 | 0.00056, 0.005379, 0.051674, 0.496387, 4.76831, 45.804544 and 440 |
| (*iii*) $S_{2,1}$ | sensitivity of S cells to drug 2 | 0.001474, 0.005429, 0.02, 0.073681, 0.271442 and 1 |
| (*iv*) $R1_{1,S}$ | sensitivity of R1 cells to drug 1 | 0, $10^{-5}$, $9.564 \times 10^{-5}$, $9.146 \times 10^{-4}$, $8.747 \times 10^{-3}$, $8.365 \times 10^{-2}$ and 0.8 |
| (*v*) $R2_{2,S}$ | sensitivity of R2 cells to drug 2 | 0, $10^{-5}$, $9.564 \times 10^{-5}$, $9.146 \times 10^{-4}$, $8.747 \times 10^{-3}$, $8.365 \times 10^{-2}$ and 0.8 |
| (*vi*) $T_1$ | mutation rate from S to R1 cells | $10^{-11}$, $2.154 \times 10^{-10}$, $4.642 \times 10^{-9}$, $10^{-7}$, $2.154 \times 10^{-6}$, $4.642 \times 10^{-5}$ and $10^{-3}$ |
| (*vii*) $T_2$ | mutation rate from S to R2 cells | $10^{-11}$, $2.154 \times 10^{-10}$, $4.642 \times 10^{-9}$, $10^{-7}$, $2.154 \times 10^{-6}$, $4.642 \times 10^{-5}$ and $10^{-3}$ |
| (*viii*) $R1_{ratio}$ | ratio of the initial R1 cell number to the total cell number | 0, $10^{-9}$, $10^{-7}$, $10^{-5}$, $10^{-3}$, $10^{-1}$, 0.9 or sampled by equation 14 |
| (*ix*) $R2_{ratio}$ | ratio of the initial R2 cell number to the total cell number | 0, $10^{-9}$, $10^{-7}$, $10^{-5}$, $10^{-3}$, $10^{-1}$, 0.9 or sampled by equation 14 |

Note: See the Model Parameters section above for detailed descriptions of the parameters.
